# Insm1 regulates the development of mTECs and immune tolerance

**DOI:** 10.1101/2023.01.14.524041

**Authors:** Wehuai Tao, Yiqiu Wei, Zhihuan Ye, Jianxue Wang, Weixin Yang, Guoxing Yu, Jieyi Xiong, Shiqi Jia

## Abstract

The *Insm1* gene encodes a zinc finger protein with known functions in neuroendocrine cells and neurons. Here we characterized the expression and function of *Insm1* in medullary thymic epithelial cells (mTECs). *Insm1* is co-expressed with Aire in majority of Insm1 or Aire positive cells, while a few Insm1 positive cells did not express Aire. Mutation of *Insm1* impair the expression of *Aire* and the generation of normal numbers of Aire-expressing mTECs during development. We detected downregulation of genes that expressed specifically in Aire-expressing mTEC and mimetic cells in *Insm1* mutant mTECs. Conversely, when *Insm1* was overexpressed in thymic epithelial cells *in vivo*, the size of the mTECs compartment was enlarged and the expression of *Aire* and genes expressed specifically in the neuroendocrine mimetic cells were increased. Mechanistically, Insm1 bound DNA in mTECs and the majority of the Insm1 binding sites were co-occupied by Aire. These Insm1 binding sites were enriched on super-enhancer regions and thus may contributed to remoted regulation. Both, mice with a thymus-specific mutation in *Insm1* or nude mice transplanted with *Insm1* mutant thymus, displayed autoimmune responses in multiple peripheral tissues. Together, our data demonstrate a role of Insm1 in development of mTECs and immune tolerance.

## Introduction

The thymus is a primary lymphoid organ where T cell progenitors undergo maturation and selection to become functional T cells. The medullary thymic epithelial cells (mTECs) control negative selection which eliminates self-antigen-recognizing T cells, and promote differentiation of regulatory T cells (Treg cells, CD4^+^Foxp3^+^CD25^+^) (1, 2). In contrast, the cortical thymic epithelial cells (cTECs) control T cell lineage commitment and positive selection. Ectopic expression of thousands of peripheral tissue-restricted antigens (TRAs) in mTECs, also called promiscuous gene expression, is believed to be a key for negative selection and the establishment of self-tolerance (3). The autoimmune regulator (Aire) is identified as the essential transcription factor that directly regulate TRAs expression and autoimmunity (4, 5). In the last two decades, studies have focused on the Aire-expressing cells and provided key molecular insights into how the TRAs expression is regulated by Aire (6–12). Particularly, Aire was identified as a chromatin looping regulator in mTECs, where Aire binds to super-enhancers and promotes the interaction between enhancers and the TRA genes (13). Studies on Aire also revealed that correct TRAs expression in the perinatal but not adult stage is essential for the establishment of self-resistance (14–16). The behind mechanism is not totally clear (17). However, studies have defined differences in the cell number of the thymic cell subpopulations during postnatal development, for instance, the perinatal cTECs (TEC progenitors) are enriched in perinatal stage and dramatically decreased with aging (17); Aire-expressing mTECs are not separated well with post-Aire mTECs in the perinatal stage, and the muscle mTECs are more enriched in perinatal than in adult stages (18). These cellular differences may have functional implications for setting tolerization during early life.

Recently, studies using single-cell RNA sequencing (scRNA-seq) identified highly differentiated mTEC subsets that bore striking similarity of molecular characteristics of the peripheral cells (19–22). These mTEC subsets are further extended by identifying the peripheral cell lineage-defining transcription factors that are required for the accumulation of these mTECs (18). The highly differentiated mTECs are named mimetic cells, which at least include FoxA-positive neuroendocrine cells, Hnf4a-positive enterohepatic cells, Sox8/SpiB-positive microfold cells, Pou2f3-positive Tuft cells, FoxJ-positive ciliated cells, Grhl-positive keratincytes and Myog-positive muscle cells (18, 23). The cellular and molecular bases of the central tolerance is thus established by TRAs expression in the Aire-expressing mTECs as well as mimetic cells under the control of additional transcription factors (23).

The mimetic cells are among the population of post-Aire mTECs (18), which is developed and differentiated from Aire-expressing mTECs (24). Therefore, nearly all mimetic cells had previously expressed Aire and thus downstream of Aire expression (18). In addition, many TRAs expressed in mimetic cells are Aire-induced genes (18). However, the differentiation of tuft cells does not depend on Aire (19)

Herzig *et al*. screened potential regulators of Aire-expression, and identified Insulinoma-associated protein 1 (Insm1) as a transcription factor expressed in mature mTECs and a candidate regulator of Aire. However, no functional analysis of Insm1 was performed in mTECs (25). We present here a genetic and molecular analysis of Insm1 in the thymus, and show that Insm1 is a novel regulator in both Aire-expressing mTECs and neuroendocrine mimetic cells.

## Results

### *Insm1* is expressed in Aire-expressing mTECs and neuroendocrine mimetic cells

To investigate the expression of *Insm1* in the thymus, we performed immunofluorescence analysis. For this we used *Insm1^+/lacZ^* and *Insm1^lacZ/lacZ^* mice in which one or two alleles of *Insm1* codon sequence were replaced by *lacZ* that encodes beta-galactosidase (β-gal) (26, 27). Therefore, the expression of *Insm1* can be monitored using β-gal. We compared signals obtained using antibodies against β-gal and Insm1, and observed that β-gal was expressed in the medulla of the thymus in both *Insm1^+/lacZ^* and *Insm1^lacZ/lacZ^* mice (Fig. S1A). Insm1 immunoreactivity overlapped with β-gal in the thymus of *Insm1^+/lacZ^* but was absent in *Insm1^lacZ/lacZ^* mice (Fig. 1A and Fig.S1A). Thus, the Insm1 antibody specific detected the endogenous Insm1 expression. Unlike the exclusive nuclear location in neuroendocrine cells (28, 29), Insm1 protein was detected in both, the nuclei and the cytoplasm of thymic cells (Fig 1B).

**Figure 1.**
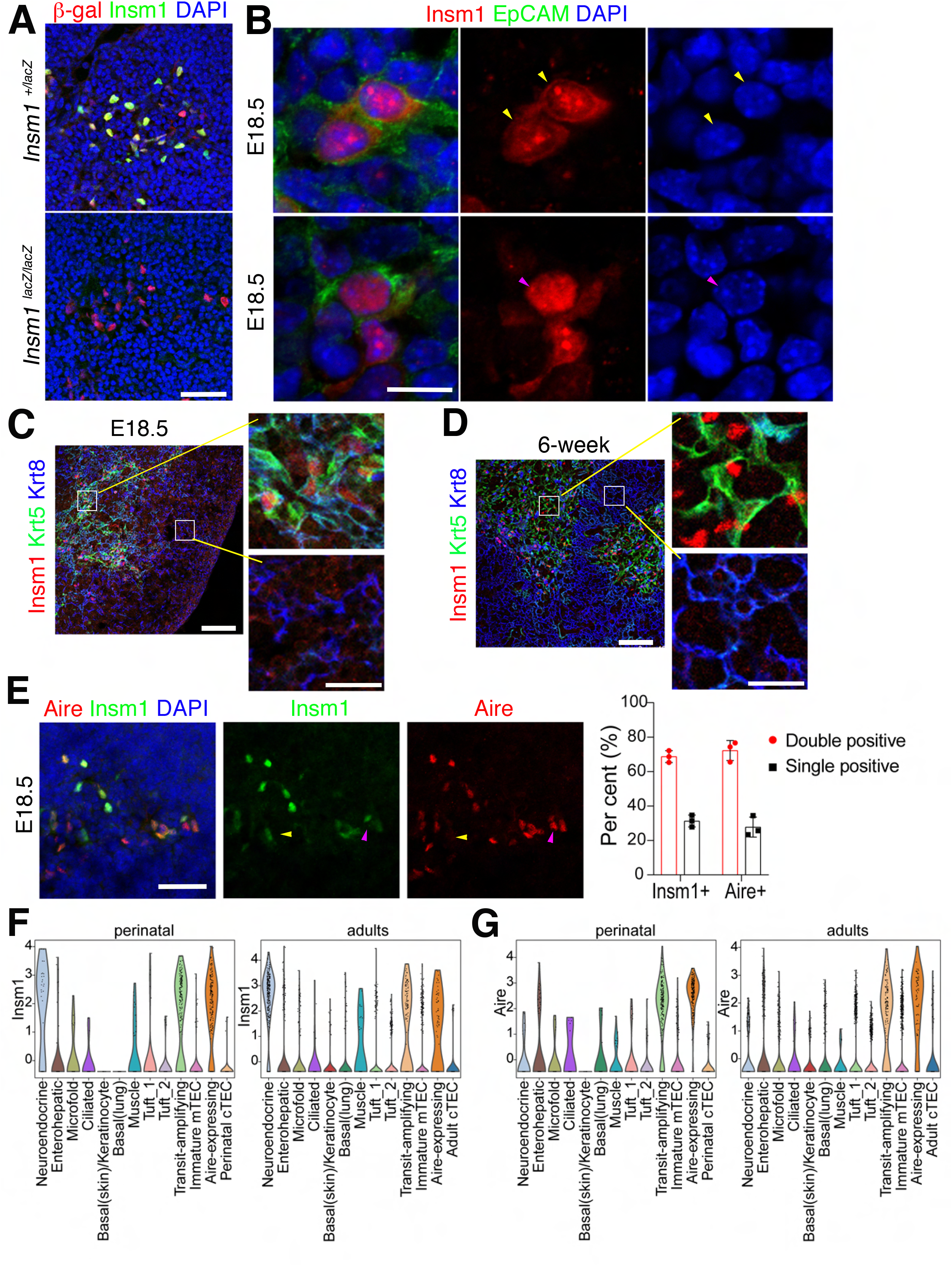
*Insm1* is expressed in thymic epithelial cells. (A) Immunofluorescence analysis using antibodies against β-gal (red) and Insm1 (green); analyzed were thymuses of E18.5 *Insm1^+/lacZ^* and *Insm1^lacZ/lacZ^* mice. DAPI (55) was used as a counterstain. Scale bar=40μm. (B) Immunofluorescence analysis of thymuses at E18.5 using antibodies against Insm1 (red), EpCAM (green) and DAPI (55). EpCAM is expressed in thymic epithelial cells and labels the cell membrane and cytoplasm. Yellow arrowheads indicated the cytoplasmic and nuclear location of Insm1, and the magenta arrowhead indicated a nucleus. Scale bar=10μm. (C,D) Immunofluorescence analysis of thymuses of E18.5 (C) and 6 weeks old (D) mice using antibodies against Insm1 (red), Krt5 (green) and Krt8 (55). Magnifications are shown in the panels on the right. Scale bar=100μm; and 20μm in magnified panels. (E) Immunofluorescence analysis detected Insm1 (green) and Aire (red) in thymuses of E18.5 mice. Insm1 is co-localized with Aire. Yellow arrowheads indicated the cell expressing only Insm1, magenta arrowheads indicated cells expressing only Aire. Scale bar=40μm. Quantification of single and double positive cells is shown on the right (animal number n=3, around 300 cells/animal/antibody were counted). (F) *Insm1* expressing in post-Aire mTECs (Pdpn^-^CD104^-^) analyzed using the published perinatal (left) and adult (right) scRNA-seq data (18). (G) *Aire* expressing in post-Aire mTECs (Pdpn^-^CD104^-^) analyzed using the published perinatal (left) and adult (right) scRNA-seq data (18).

The thymic epithelial compartment is formed by cTECs and mTECs, and contains also lymphocytes and dendritic cells (30). To define the cell-type that expresses *Insm1*, we performed immunofluorescence using antibodies against Insm1 and cell type specific markers in both, the fetal (E18.5) and adult thymus. We did not observe co-expression of Insm1 with the lymphocyte specific marker CD45, or the dendritic cell marker Cd11b and Cd11c in thymuses of E18.5 and 6-week old mice (Fig. S1B,C). However, Insm1 was co-expressed with the mTECs marker keratin 5 (Krt5) but not the cTECs marker keratin 8 (Krt8) (Fig. 1C,D). Using Insm1/Aire double antibodies staining, we further observed that 70% of the Insm1-positive cells co-expressed Aire and vice versa (Fig. 1E). The overlapping expression pattern of Insm1 and Aire was also observed in thymuses of adult mice (Fig S1D). Moreover, the Aire-dependent TRAs insulin and somatostatin were co-expressing with Insm1 in the thymus of adults (Fig S1E and S2A). To investigate the expression of *Insm1* in post-Aire populations, we used the published single cell RNA-seq data, which were generated from the isolated post-Aire mTECs using the two makers that specifically downregulated in post-Aire mTECs, podoplanin (Pdpn) and integrin β4 (CD104), i.e., the Pdpn^-^ CD104^-^ cells separated from CD45^-^EpCAM^+^MHCII^lo^Ly51^-^ mTECs (18). The scRNA-seq data showed that the isolated cells contained mostly the post-Aire mTECs, but also cells that are not fully separated from post-Aire mTECs which include immature mTECs, transit-amplifying mTECs (the direct progenitor of Aire-expressing mTECs), Aire-expressing mTECs, and cTECs (18) (Fig S2B). We found that *Insm1* was expressed in neuroendocrine cells in the post-Aire mimetic cells in both perinatal and adult stages (Fig 1F and Fig S2B-D), while Aire expressing was not appeared in the mimetic cells (Fig 1G and Fig S2B-D). We further verified the co-expressing of Insm1 with neuroendocrine mimetic cell markers Foxa2 and chromogranin A in adult thymus (Fig 2A,B). In summary, *Insm1* is expressed in Aire-expressing mTECs and post-Aire neuroendocrine mimetic cells.

**Figure 2.**
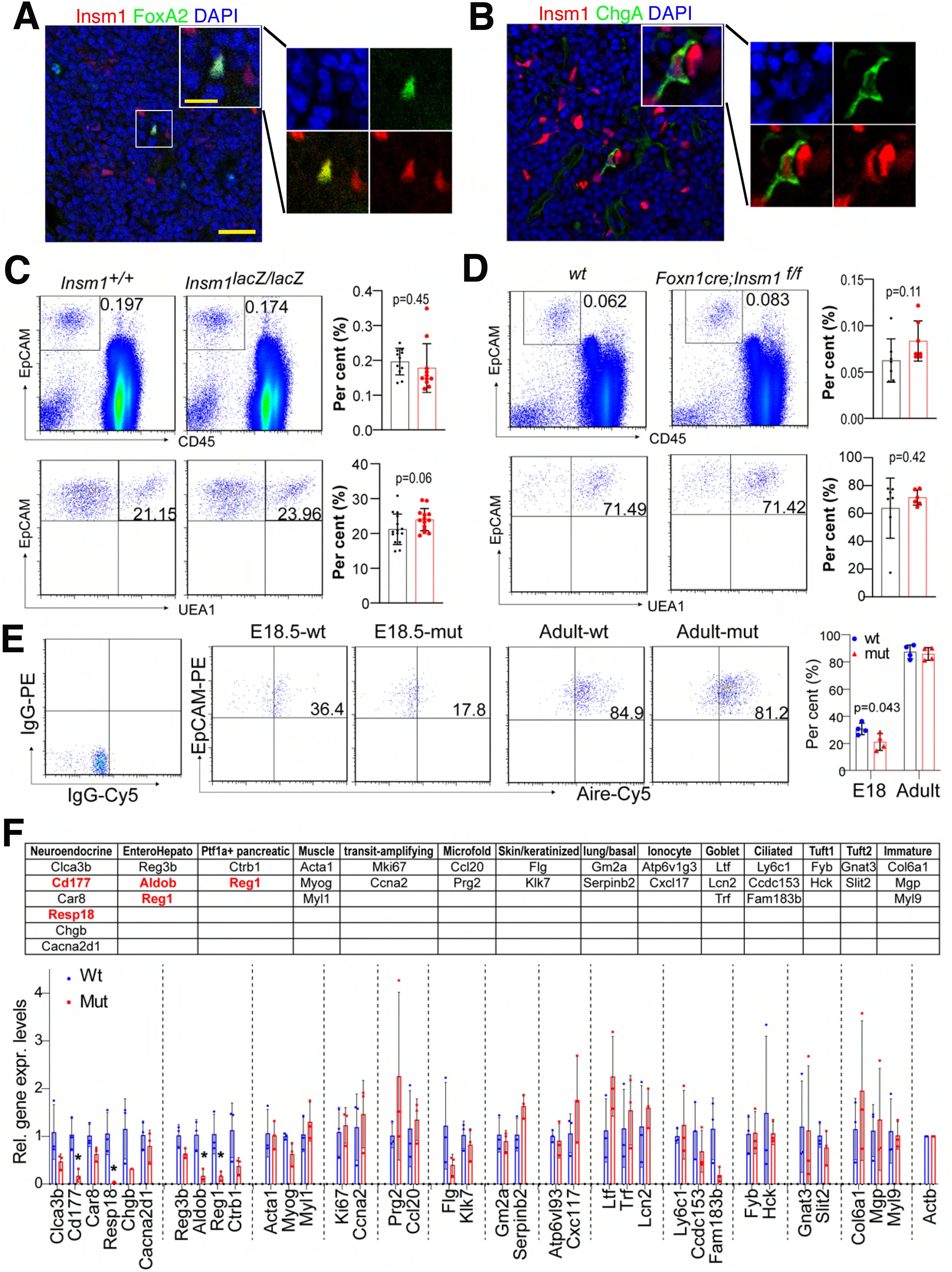
Cellular phenotypes observed in the *Insm1* mutant thymus. (A,B) Immunofluorescence analysis of thymuses of 6-week-old mice using antibodies against (A) Foxa2 (green) and Insm1 (red), and (B) ChgA (green) and Insm1 (red). Scale bar=25μm and 10μm in the main and magnified panels, respectively. (C) Flow cytometry analysis of CD45^-^ /EpCAM^+^ epithelial cells (the upper panel) and CD45^-^/EpCAM^+^/UEA1^+^ mTECs (the lower panel) in thymuses of *Insm1* mutant and wildtype mice (n=11). (D) Flow cytometry analysis of CD45^-^/EpCAM^+^ epithelial cells (the upper panel) and CD45^-^/EpCAM^+^/UEA1^+^ mTECs (the lower panel) in thymuses of *Thy-cKO* and wildtype mice (n=6). (E) Flow cytometry analysis of Aire^+^ cells in *Insm1* mutant and wildtype control animals at the age of E18.5 and adult (animal number n=4 for each genotype). Isotype control staining for Aire was showed on the left, quantifications were showed on the right of the panel. (F) Quantitative RT-PCR analysis of the expression of the mimetic cells specific genes. Upper panel showed the genes specific expressed in the mimetic cell types refereed to the published data (18). Red labeled the downregulated genes in *Insm1* mutants. Lower panel showed the qRT-PCR results. Significant downregulated genes were labeled red in the upper panel. Data in the figure panel are presented as means ± SD, statistical significance was assessed by 2-tailed unpaired Student’s t-test. Statistically significant was defined as *p*<0.05.

### Tissue morphology and cell populations in *Insm1* mutant thymus

There was no difference in the appearance of size and shapes of thymus at E18.5 in mutants (Fig S2D). The distribution and relative amounts of mTECs and cTECs identified by Krt5 and Krt8, respectively, were comparable between wildtype and *Insm1* mutant mice (Fig S2E,F). We used flow cytometry to further investigated cell populations of mTECs. The number of EpCAM labeled thymic epithelial cells (CD45^-^EpCAM^+^) and UEA1 labeled mTECs (CD45^-^EpCAM^+^UEA1^+^) were comparable in wildtype controls and *Insm1* mutants both at E18.5 and adults (Fig.2C,D). However, the proportions of Aire-positive cells were significantly decreased in mTECs (CD45^-^ EpCAM^+^Ly51^-^) of *Insm1* mutants at E18.5 but not altered in adult animals (Fig.2E). Thus, the *Insm1* mutation affected the development of Aire-positive mTECs.

Using qRT-PCR, we examined the expression of genes that were identified to be specifically expressed in each of the mimetic cells (18) (Fig 2F upper panel). We found that genes specifically expressed in neuroendocrine, enterohepatic and Ptf1a^+^pancreatic cells were downregulated in *Insm1* mutant adult mice (Fig 2F lower panel). The data indicated that the mimetic cells, i.e., neuroendocrine, enterohepatic and Ptf1a^+^pancreatic mimetic cells, were affected by the *Insm1* mutation.

### Decreased expression of *Aire* and mTEC genes in *Insm1* mutants

As we observed a decreased number of Aire-positive cells in the developing *Insm1* mutant thymus, we investigated the expression levels of Aire. The immunofluorescence intensity which indicated the Aire protein level was moderately but significantly decreased in mTECs of *Insm1* mutant mice at E18.5 (Fig.3A). Using real-time quantitative reverse transcription PCR (qRT-PCR), we detected decreased transcript levels of *Aire* in mTECs in both, *Insm1* mutant E18.5 mice and adult mice with thymic specific mutation in *Insm1 (Foxn1Cre;Insm1^flox/flox^*, subsequently called *Thy-cKO*) (Fig. 3B,C). Thus, the *Insm1* mutation resulted in decreased expression of *Aire*.

**Figure 3.**
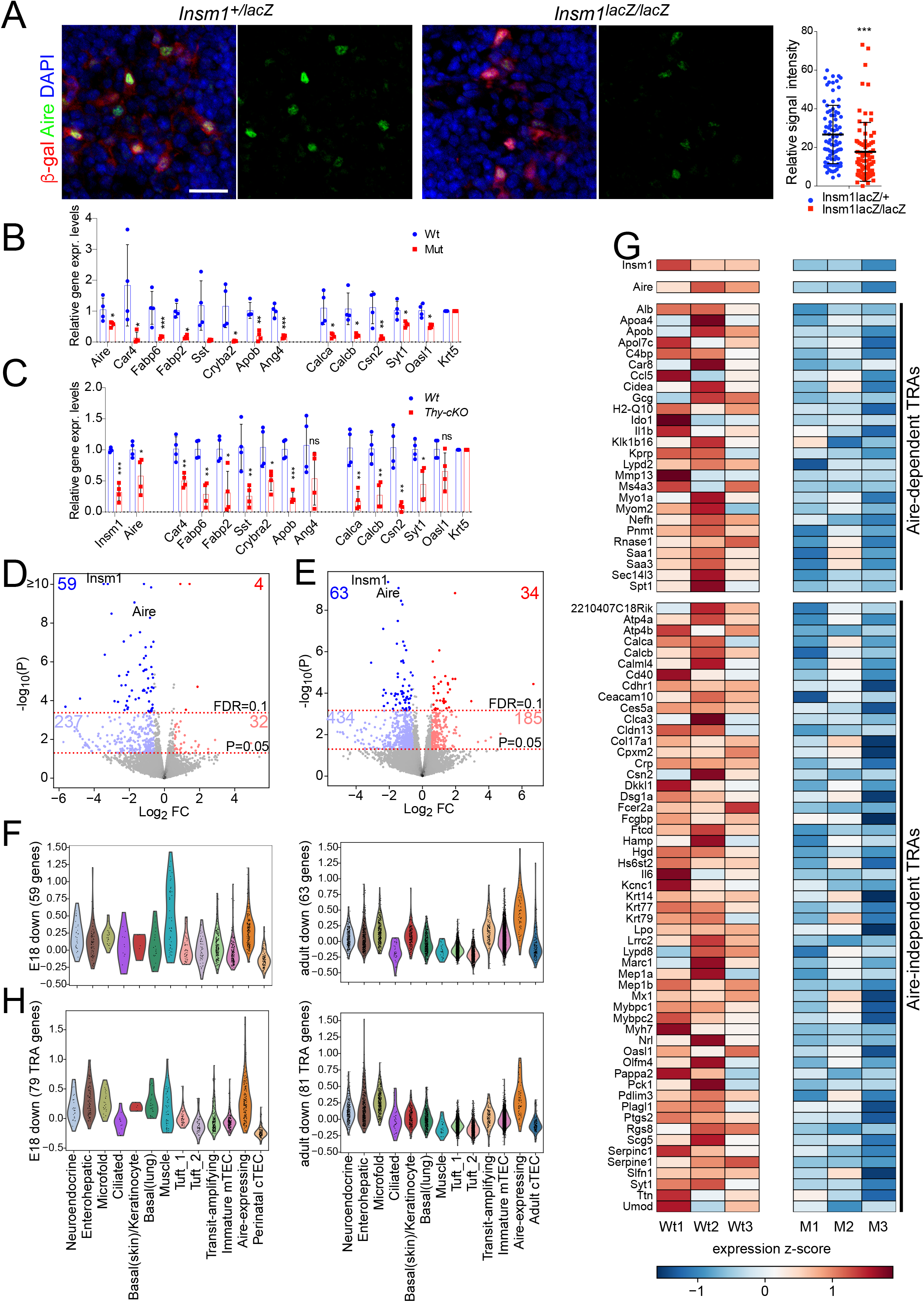
Changes in gene expression in the *Insm1* mutant thymus. (A) Immunofluorescence analysis of the expression of β-galactosidase (β-gal, red) and Aire (green) in thymuses of E18.5 mice. DAPI (55) was used as a counterstain. Quantifications of the fluorescence intensity are shown on the right (animal number n=4 for each genotype, 80-100 Aire+/β-gal+ cells were counted). Scale bar=20μm. (B,C) Quantitative RT-PCR analysis of the expression of *Aire* and TRA genes in mTECs of E18.5 (B) and at adults (C) (n=4). Data in the figure panel are presented as means ± SD, statistical significance was assessed by 2-tailed unpaired Student’s t-test. ns: *p*>0.05; *: *p*<0.05, **: *p*<0.01, ***: *p*<0.001. (D,E) Volcano blots of the RNA-seq data in *Insm1* mutants versus wildtype controls at E18.5 (D) and adult stages (E). FDR≤0.1 or *p*-value≤0.05 combined with Fold change≥1.5 were used for identifying the dysregulated genes. The numbers of the dysregulated genes were showed in figures accordingly. (F) Expression distribution of the Insm1-dependent genes (FDR≤0.1,FC≥1.5) in mimetic cells. Presented by the violin plots of the gene score for each cell of the post-Aire mTECs. Left panel, downregulated gene in E18.5 *Insm1* mutant mice plotted on published perinatal scRNA-seq data; right panel, downregulated genes in adult *Insm1* mutant mice plotted on published adult scRNA-seq data. (G) Heatmap of dysregulated TRA genes expressed in mTEC of *cKO* mice. (H) expression distribution of the Insm1-dependent TRA genes (p≤0.05, FC≥1.5) in mimetic cells. Presented by the violin plots of the gene score for each cell of the post-Aire mTECs. Left panel, the dysregulated TRAs genes in E18.5 *Insm1* mutant mice plotted on published perinatal scRNA-seq data; right panel, the dysregulated TRAs genes in adult *Insm1* mutant mice plotted on published adult scRNA-seq data.

To analyze the global gene expression changes in *Insm1* mutant thymus, we performed RNA-seq using mTECs isolated from both *Insm1* mutant E18.5 and *thy-cKO* adult mice. To our surprise, differential gene expression test by DESeq2 under cutoff of FDR≤0.1 and fold change≥1.5, identified 63 and 97 dysregulated genes in E18.5 and adult mice, respectively (Fig 3D,E and Table S1,S2). The numbers of the dysregulated genes were far less than that of Aire-regulated genes reported in a previous study (31). Nevertheless, *Aire* expression was detected downregulated in both E18.5 and adults in the RNA-seq data (Fig 3D,E), which was consistent with the observation in qRT-PCR and immunofluorescence analysis (Fig 3A-C). To characterize the *Insm1* effects on different mimetic cell types, we analyzed the expression pattern of the Insm1-dependent genes in the published scRNA-seq data that were generated from post-Aire mTECs (18). The downregulated genes enriched in several types of mimetic cells as well as in the unseparated Aire-expressing mTECs both in E18.5 (Fig 3F left) and in adult (Fig 3F right). Thus, although moderate numbers of the dysregulated genes were identified in *Insm1* mutants, Insm1 regulates the gene expression in Aire-expressing mTECs and mimetic cells.

Considering that each of the TRAs is typically expressed in only 1-5% of mTECs (32), we used a loose cutoff (*p*-value ≤0.05 and fold change≥1.5) to define trends of TRAs expression(Fig 3D,E), which distinguished 78 and 81 downregulated TRA genes in the mTECs of E18.5 and adult animals, respectively (Fig S3A, Fig 3G and Table S3,S4). Using Qrt-PCR, we verified the altered expression of a subset of these dysregulated genes (Fig 3B,C). We compared the downregulated genes observed in *Insm1* mutant mTECs with the ones identified in *Aire* mutant mTECs (31). Among the dysregulated TRA genes, we found both, Aire-dependent and -independent TRAs, to be downregulated (Fig 3G and Fig S3A). Among the 81 downregulated TRA genes in adults, twenty-six was downregulated in both, *Aire* and *Insm1* mutants. However, fifty-five (68%) of the TRA genes were unique to *Insm1* mutant mTECs (Fig.3G). Thus, Insm1 regulates the expression of TRA genes in mTECs, and effects both, Aire-dependent and -independent TRAs.

Using the published scRNA-seq data (18), we assigned the downregulated TRAs to the mimetic cell types. Among the mimetic cell types, *Insm1* expression is restricted in neuroendocrine cells (Fig 1F). However, the downregulated TRAs were not restricted to neuroendocrine mimetic cells. Instead, these TRAs was found in multiple mimetic cell types, i.e., the enterohepatic and microfold cells, in both E18.5 and adult animals (Fig 3H). We investigated the tissue types where TRA genes are dominantly expressed using public microarray data (33). We noticed that Insm1-dependent TRAs were expressed in all the tested peripheral tissues and with relative higher frequency in the gastrointestinal tract and neuroendocrine tissues (Fig.S3B).

### Insm1 promotes *Aire* and TRA expression

We induced over-/ectopic-expression of *Insm1 (Insm1OE) in vivo* using genetic tools (Fig S4), i.e., a mouse line that harbors a *loxp-STOP-loxp-Insm1* cassette in the *Rosa26* locus, and the *Foxn1Cre* transgenic allele that allows thymus specific expression. We observed increased numbers of Insm1-positive cells and enlarged Krt5-positive mTECs areas in the *Insm1OE* thymus (Fig 4A). Moreover, the Aire-positive cells were increased in the thymic sections, although the increase of such cells was less obvious as the one of Insm1-positive cells (Fig 4A). Increased expression levels of *Aire* were detected by qRT-PCR using RNA isolated from the whole thymus or from mTECs (Fig 4B,C). Interestingly, four of the nine TRA genes that were downregulated in *Insm1* mutant thymuses showed significantly increased expression upon *Insm1* overexpression (Fig 4B,C). We investigated the expression of mimetic cell marker genes that was downregulated in *Insm1* mutant mTEC. *Insm1* overexpressing promoted the neuroendocrine cells specific marker genes expression (Fig 4B,C). Thus, Insm1 promoted mTECs fate and the expression mTEC specific genes such as *Aire*, TRAs and neuroendocrine-specific markers.

**Figure 4.**
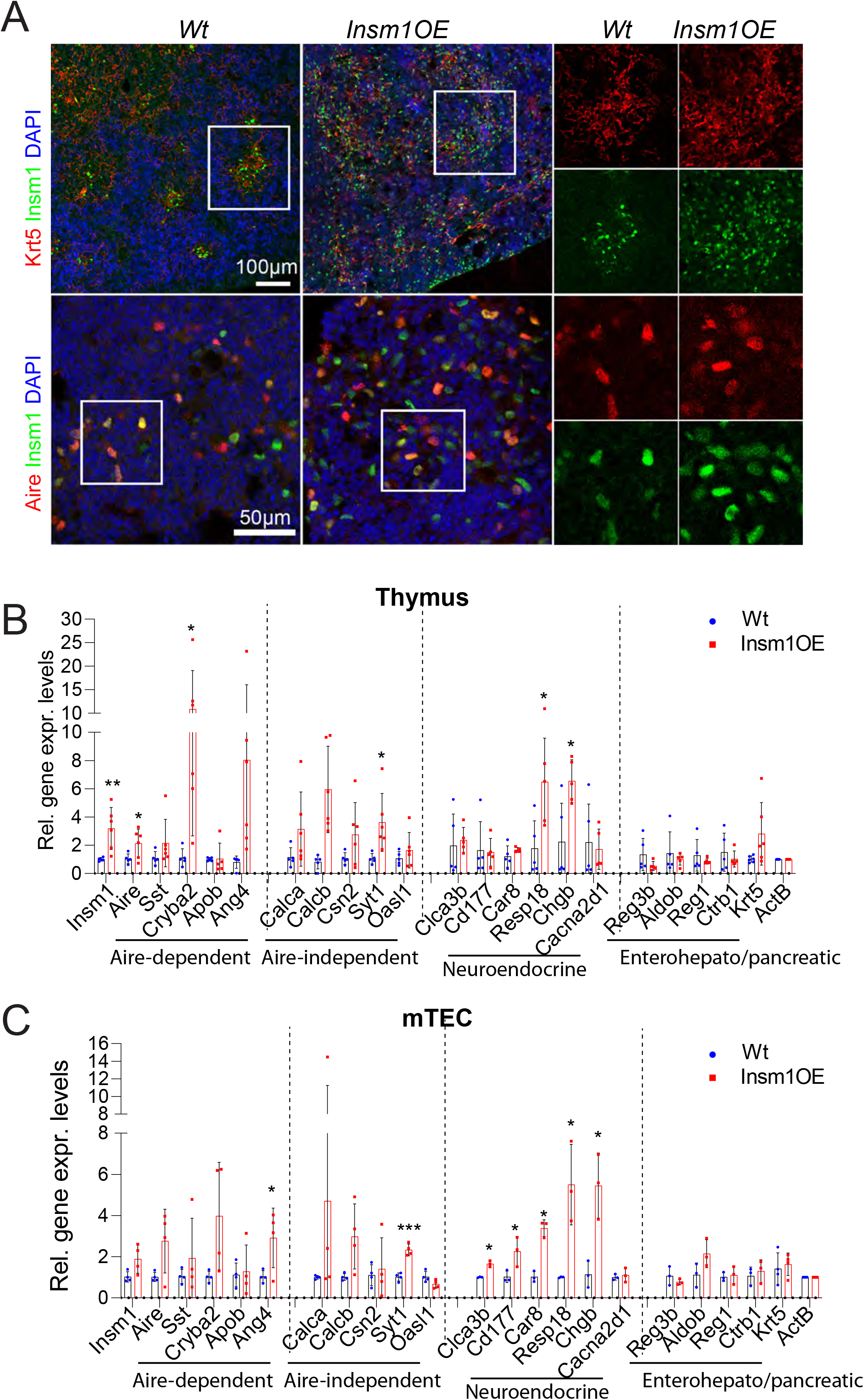
Insm1 promotes thymic gene expression in the thymus. (A) Immunofluorescence analysis of the expression of Insm1 (green), and Krt5 (red, upper panels), Aire (red, lower panels) in *Insm1*-overexpression (*Insm1OE*) thymuses of postnatal 2-day animals. DAPI (55) was used as a counterstain. (B-C) Comparison of gene expression in *Insm1OE* and control thymuses using postnatal 2-day animals (animal number n=5-6) (B) and mTECs (animal number n=3-4) (C) using qRT-PCR. Data are presented as means ± SD, statistical significance was assessed by 2-tailed unpaired Student’s t-test. *: *p*<0.05, **: *p*<0.01, ***: *p*<0.001.

#### Insm1 binds on promoters and super-enhancers in mTECs

Insm1 was identified as a transcriptional factor in endocrine cells (28). To investigate the mechanism of Insm1 function, we examined the genome wide DNA-binding of Insm1 using CUT & Tag analysis. We detected 5,206 Insm1 binding peaks (union of 3 repeats, q<1e-5) in mTEC of E18.5 and 1,458 (union of 2 repeats, q<1e-5) peaks in mTEC of adults (Fig 5A,B). Although a less numbers of Insm1 binding peaks were identified in adults, 1,382 (94.7%) peaks identified in adult mTEC were overlapped with the peaks detected in E18.5 mTEC, indicating a significant amount of consensus Insm1 binding on DNA in mTEC during development.

**Figure 5.**
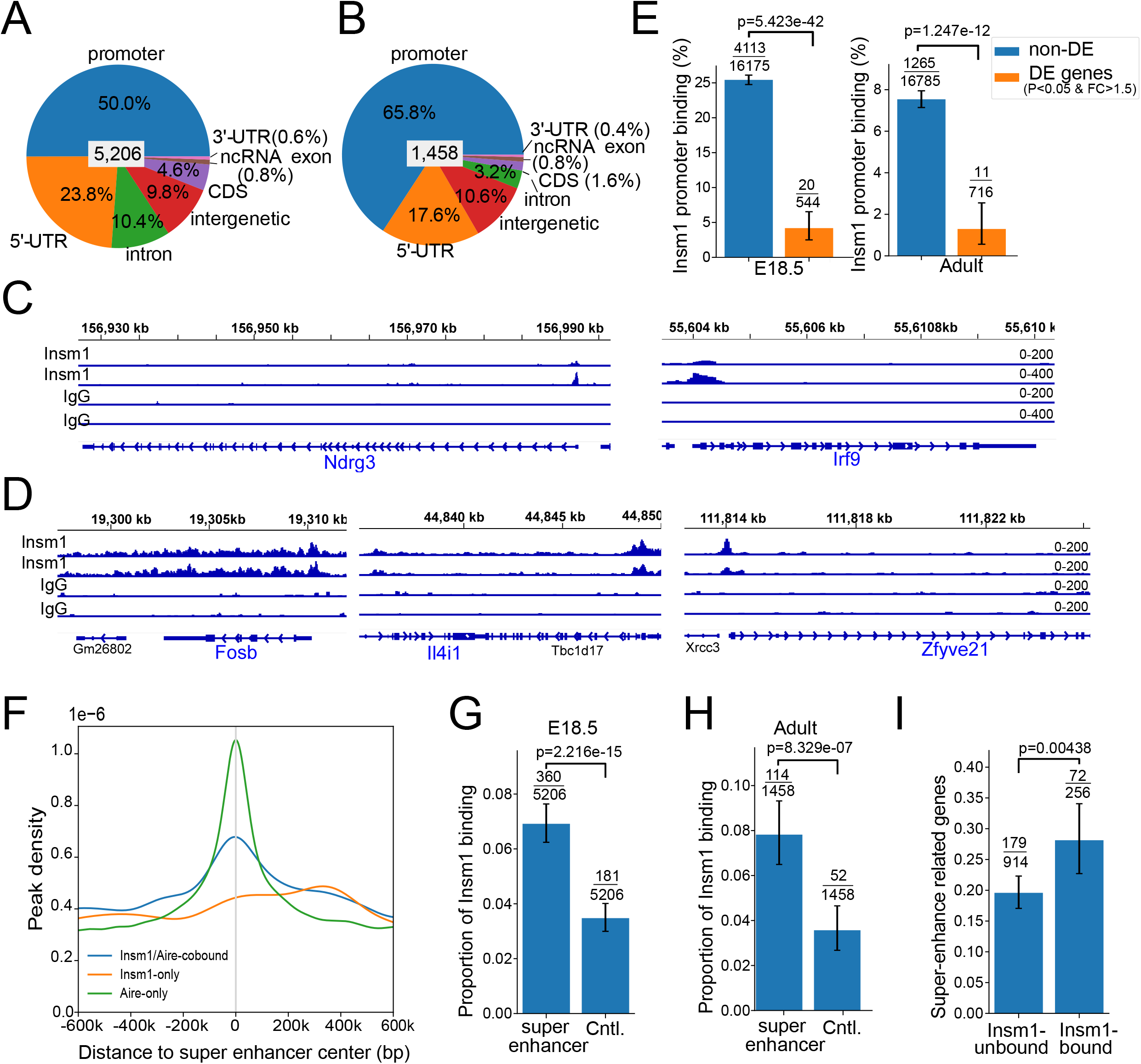
Insm1 binds to super-enhancer loci. (A, B) Distribution of Insm1 binding sites in mTEC genome at the E18.5 stage (A) and at the adult stage (B). (C,D) the Insm1 binding traces on dysregulated genes at the E18.5 stage (C) and at the adult stage (D). (E) Proportions of genes with Insm1 binding sites in the promoter regions (−2000bp to +500bp of TSS) at the E18.5 stage (left) and adult stage (right). Insm1 binding sites are significantly depleted on the promoters of dysregulated genes (F) Density curves of detected loci Inms1 and Aire co-binding (55), Insm1-only binding (orange), and Aire-only binding (green), aligned to the center (0 position) of the super-enhancers. (G,H) Proportion of Insm1 binding peaks overlapping with super-enhancer and control loci at the E18.5 stage (G) and the adult stage (H). An identical length sequence located 200kb plus the length of super-enhancer away from each super-enhancer was selected as the control sequence. (I) Proportions of super-enhancers located within ±500kb of genes that were dysregulated (*p*-value≤0.05, FC≥1.5) in *Insm1* mutant mTECs at the E18.5 stage. Stratified by the super-enhancers bound or unbound by Insm1. In (E), (G), (H) and (I), numerators and denominators of each proportion are given above the bars. The bars show 95% confidential intervals. The *p*-values of Fisher test are also given.

Although the majority of Insm1 binding sites were located on promoter regions in both E18.5 (50%) and adults (65.8%) (Fig 5A,B), only two (Fig 5C, E18.5) and three (Fig 5D, adult) of these Insm1 bound genes were dysregulated in Insm1 mutation (Fig 5C,D). Furthermore, we found the dysregulated genes, which were identified by a looser cutoff (*p*-value<0.05, FC>1.5) in the RNA-seq data, were not enriched but depleted for Insm1 binding on the promoter (Fisher test, *p*<0.0001) (Fig 5E). These evidences suggest that Insm1 binds on promoters but did not significantly contribute to the correlated gene expression.

Insm1 is co-expressed with Aire in mTECs (Fig1E, Fig S1D), we therefore compared the DNA binding of Insm1 with that of Aire identified in published data sets (31, 34). The majority of Insm1 binding sites (72-78%) were co-occupied by Aire (Fig S5A,B). Since Aire was reported binding on super-enhancers and performing regulatory roles in mTEC (13), we investigated the Insm1 binding sites and found significantly enriched binding of Insm1 on super-enhancers (Fig.5F-H). We further found that the dysregulated genes were significantly over-represented within 500kb of the Insm1-binding super-enhancers at E18.5 mTECs (Fig. 5I). These results imply that Insm1 could bind on super-enhancers and participate in the gene expression regulation in the developing mTECs.

### Autoimmune responses in nude mice transplanted with *Insm1* mutant thymuses

We used thymus transplantation in nude mice to study autoimmune reactions. For this, T cells depleted thymuses that were isolated from fetal control (*Insm1^+/lacZ^*) and mutant (*Insm1^lacZ/lacZ^*) mice were transplanted into the renal capsule of six-week old nude mice (35) (Fig.6A,B). Eight weeks after transplantation, the size and structure of the transplanted thymuses were similar regardless whether they derived from control or *Insm1* mutant animals (Fig.6C and Fig.S6). Further, the numbers of CD4+ and CD8+ cells detected by flow cytometry were comparable in the transplanted control and mutant thymuses (Fig.6D). However, the spleens of nude mice transplanted with a mutant thymus (*KO/nu* mice) were increased in size and weight compared to the one of animals that received a control transplant (*Het/nu* mice) (Fig. 6E). Furthermore, the structure of the spleen was altered (Fig 6F), but the lymph nodes were unchanged (Fig S6). In particular, spleen follicles of *KO/nu* mice were smaller than that those of *Het/nu* mice (Fig. 6F). We investigated lymphocyte infiltration of various organs using hematoxylin eosin (H&E) staining, that detects the accumulation of lymphocyte nuclei. Increased infiltration was observed in pancreatic islets, lungs, kidneys and salivary glands of *KO/nu* mice (Fig.6G,H), but not in other investigated tissues (Fig.S6).

**Figure 6.**
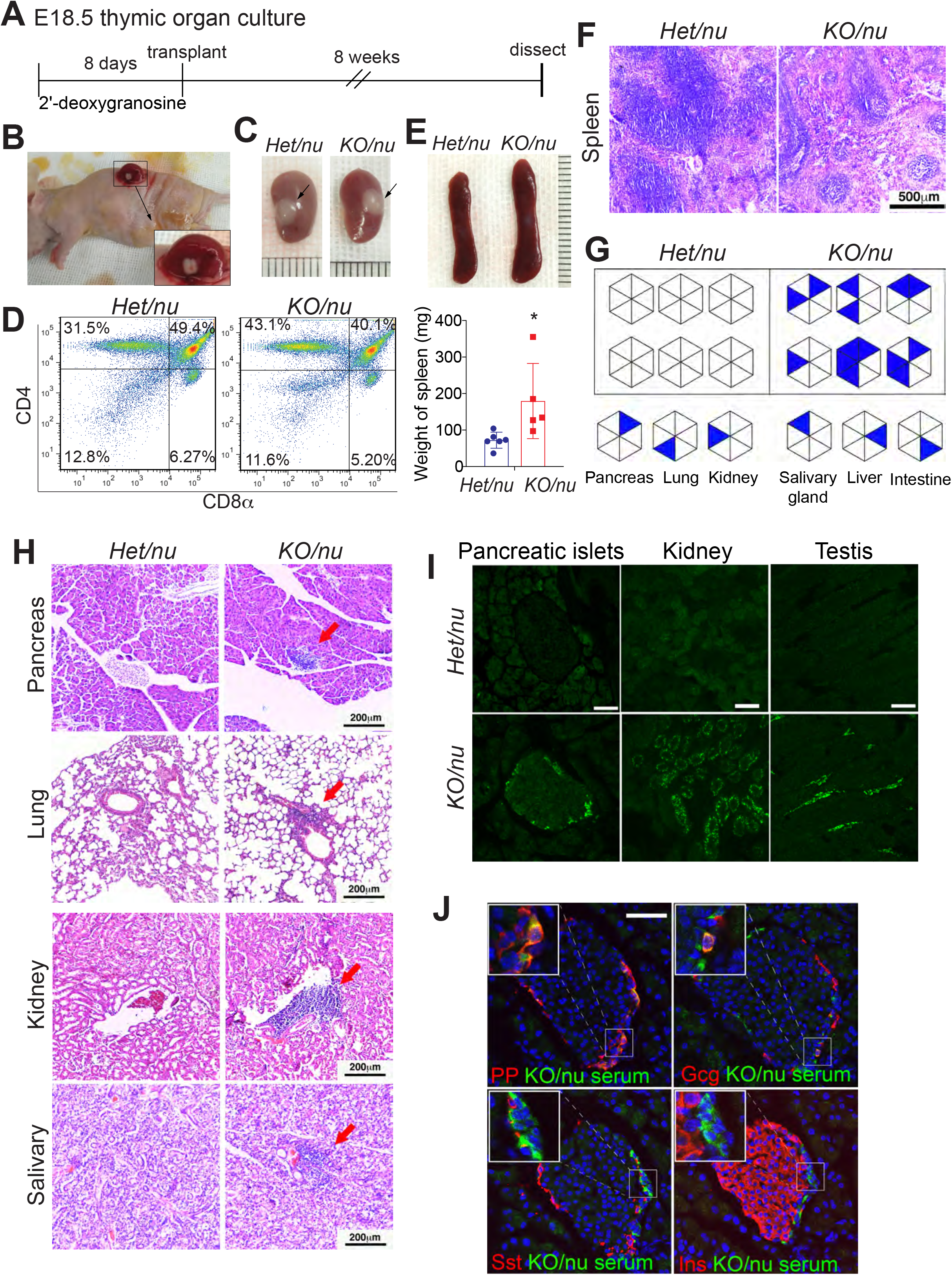
Transplantation of *Insm1* mutant thymuses in nude mice. (A) Schematic outline for transplantation experiments. Thymuses were isolated, depleted for lymphocytes by 8-day culture in 2-deoxygranosine, transplanted into nude mice, and the mice were analyzed 8 weeks after transplantation. (B) Transplantation of the thymus under the kidney capsule of the nude mouse. Enlarged picture shows the kidney with the transplanted thymus. (C) Isolated kidneys showing the thymuses under the kidney capsule 8-week after transplantation. *Het/nu*: the thymus was isolated from an *Insm1^+/lacZ^* mouse and transplanted into a nude mouse; *KO/nu*: the thymus was isolated from an *Insm1^lacZ/lacZ^* mouse and transplanted into a nude mouse. (D) CD4 and CD8α staining and flow cytometry analysis of thymocytes isolated from the transplanted thymuses of *Het/nu* and *KO/nu* mice (n=3). (E) Appearance (upper panel) and quantification of the weight (lower panel) of spleens isolated from *Het/nu* and *KO/nu* mice (n=5-6). (F) H&E staining shows the structures of spleens isolated from *Het/nu* and *KO/nu* mice. (G) Summary of lymphocytes infiltration in multiple tissues of *Het/nu* and *KO/nu* mice (n=6). (H) H&E staining of the pancreas, the lung, the kidney and the salivary gland isolated from *Het/nu* and *KO/nu* mice. 6-8 sections from the non-serial sections were used for H&E staining for each tissue. Red arrows indicate sites of lymphocytes infiltration. (I) Immunostaining using serum (green) isolated from *Het/nu* and *KO/nu* mice on sections that were prepared from *Rag1^-/-^* mice. 3-4 sections collected from the non-serial sections of two *Rag1^-/-^* mice were used for the serum staining for each tissue. (J) Shown is the co-staining of the *KO/nu* serum (green) with pancreatic islet specific hormones PP, Gcg, Sst and Insulin (red, as indicated in the panels). Statistical data are presented as means ± SD, significance was assessed by 2-tailed unpaired Student’s t-test. *: *p*<0.05.

Next, we analyzed autoimmune reactions by the detection of autoimmune antibodies. Among the 13 investigated tissues, we observed autoimmune antibody reactions in pancreatic islets, kidney, and testis by staining these tissues with the serum obtained from the *KO/nu* mice (Fig 6I and Fig S7). In particular, serum of *KO/nu* mice detected subsets of pancreatic polypeptide (Pp) positive cells, glucagon (Gcg) positive alpha cells, and somatostatin (Sst) positive delta cells, whereas insulin (Ins) positive beta cells were rarely detected (Fig 6J). Thus, autoimmune antibody against PP-, alpha- and delta-cells were present in *KO/nu* mice. To sum up, the transplantation of *Insm1* mutant thymuses into nude mice results in autoimmune responses in multiple tissues.

### Thymus specific *Insm1* mutant mice show autoimmune responses

We employed thymus-specific *Insm1* mutant mice, *Foxn1Cre;Insm1^flox/flox^* (*Thy-cKO*), and littermates *Foxn1Cre* or *Insm1^flox/flox^* controls to further examine the autoimmune phenotype. The *Thy-cKO* mice were born healthy and displayed no obvious abnormalities in adulthood. We observed decreased expression levels of *Insm1, Aire* and the Aire-dependent and - independent TRAs in mTECs of *Thy-cKO* mice (Fig 3C), which is similarly to the changes observed in *Insm1* null mutant mTECs (Fig 3B). CD4^+^ and CD8^+^ T cells was properly generated in the thymus (Fig 7A) and developed in lymph nodes (Fig 7B) of *Thy-cKO* mice, as analyzed by flow cytometry. However, we detected significantly decreased numbers of Treg cells (CD8^-^ CD4^+^Foxp3^+^CD25^+^) in the thymus of *Thy-cKO* mice (Fig 7C), and we observed a trend towards a reduced number of Treg cells in lymph nodes that did not reach statistical significance (Fig 7D).

**Figure 7.**
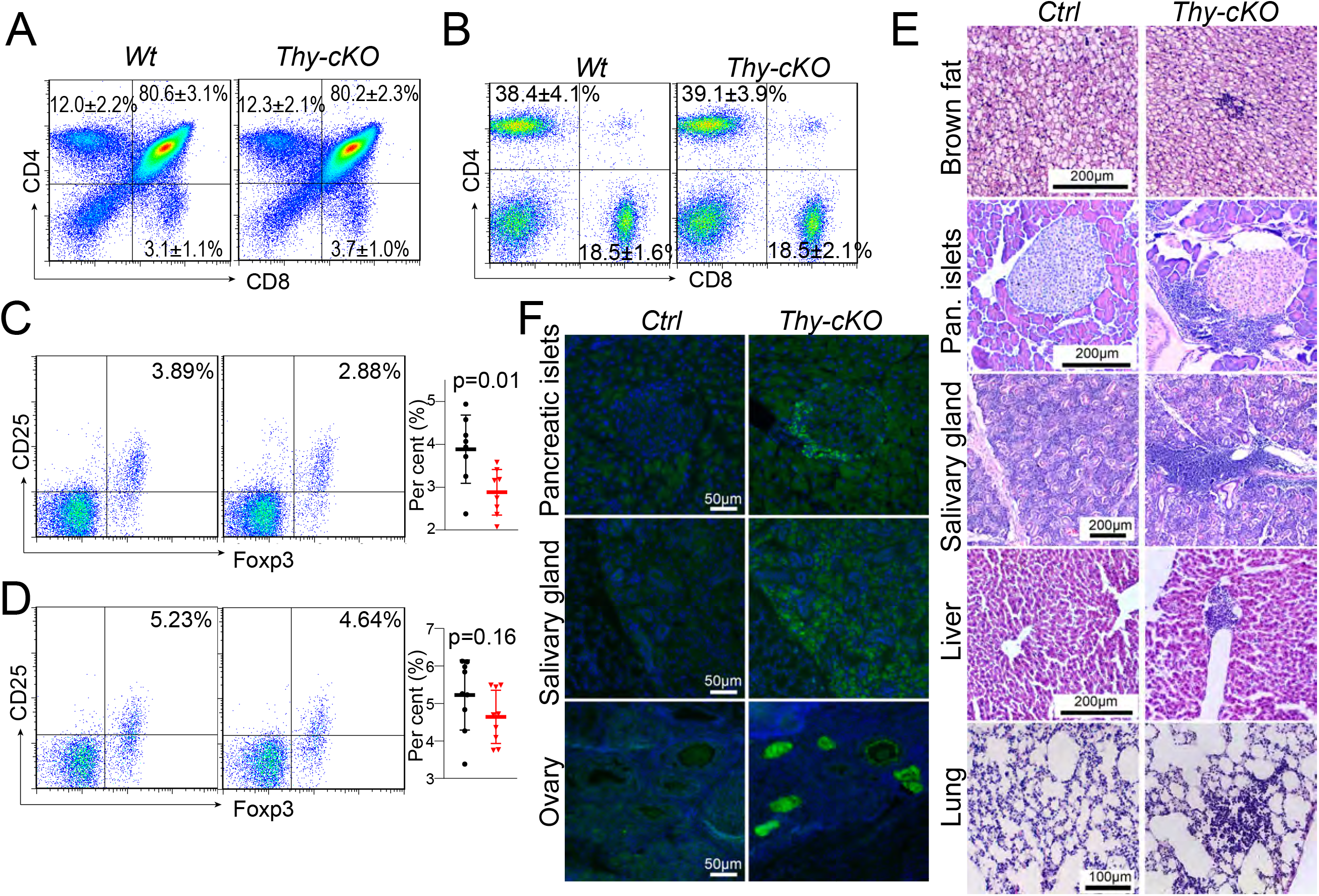
Autoimmune phenotype in thymus specific *Insm1* mutant mice. (A) CD4 and CD8α staining and flow cytometry analysis of thymocytes from thymuses of *Thy-cKO* and wildtype mice (mean and SD showed in figures, n=10-12). (B) CD4 and CD8α staining and flow cytometry analysis of thymocytes from axillary lymph nodes of *Thy-cKO* and wildtype mice (mean and SD showed in figures,n=9). (C, D) Flow cytometry analysis of CD4^+^/CD25^+^/Foxp3^+^ Treg cells from the thymus (C, n=8) and the axillary lymph node (D, n=9) of *Thy-cKO* and wildtype mice. (E) H&E staining of brown fat, pancreas, salivary gland, liver and lung in *Thy-cKO* and *Insm1^f/f^* or *Foxn1Cre* control mice. 6-8 sections from the non-serial sections were used for H&E staining for each tissue. Animal numbers were list in Table 1. (F) Immunostaining using serum isolated from *Thy-Cko* and *Insm1^f/f^* or *Foxn1Cre* mice on sections of pancreas, salivary gland and ovary prepared from *Rag1^-/-^* mice. 3-4 sections collected from the non-serial sections of two *Rag1^-/-^* mice were used for the serum staining for each tissue. The animals that the serum was collected were list in Table 2. Statistical data are presented as means ± SD, significance was assessed by 2-tailed unpaired Student’s t-test. *P*>0.05 in figure A and B, *p*-values showed in figures C and D.

We investigated the infiltration of lymphocytes into various tissues at 6-week, 6-, 12- and 18-month-old *Thy-cKO* animals. Generally, we observed pronounced lymphocyte infiltration in multiple organs that increased in frequency with the age of the animals. Infiltrated lymphocytes were observed in salivary glands of 6-week-old and in multiple organs of 1.5-year-old *Thy-cKO* mice, the infiltrated organs including the pancreas, the slavery gland, the liver, the lung and the brown fat (Table 1 and Fig 7E). Further, we detected autoimmune antibodies against multiple tissues including the pancreas, the salivary gland and the ovary when serum of *Thy-cKO* mice was used for immunostaining (Table 2 and Fig 7F). Thus, mutation of *Insm1* in the thymus resulted in autoimmune responses in multiple periphery organs in a subset of the *thy-cKO* mice.

**Table 1.**
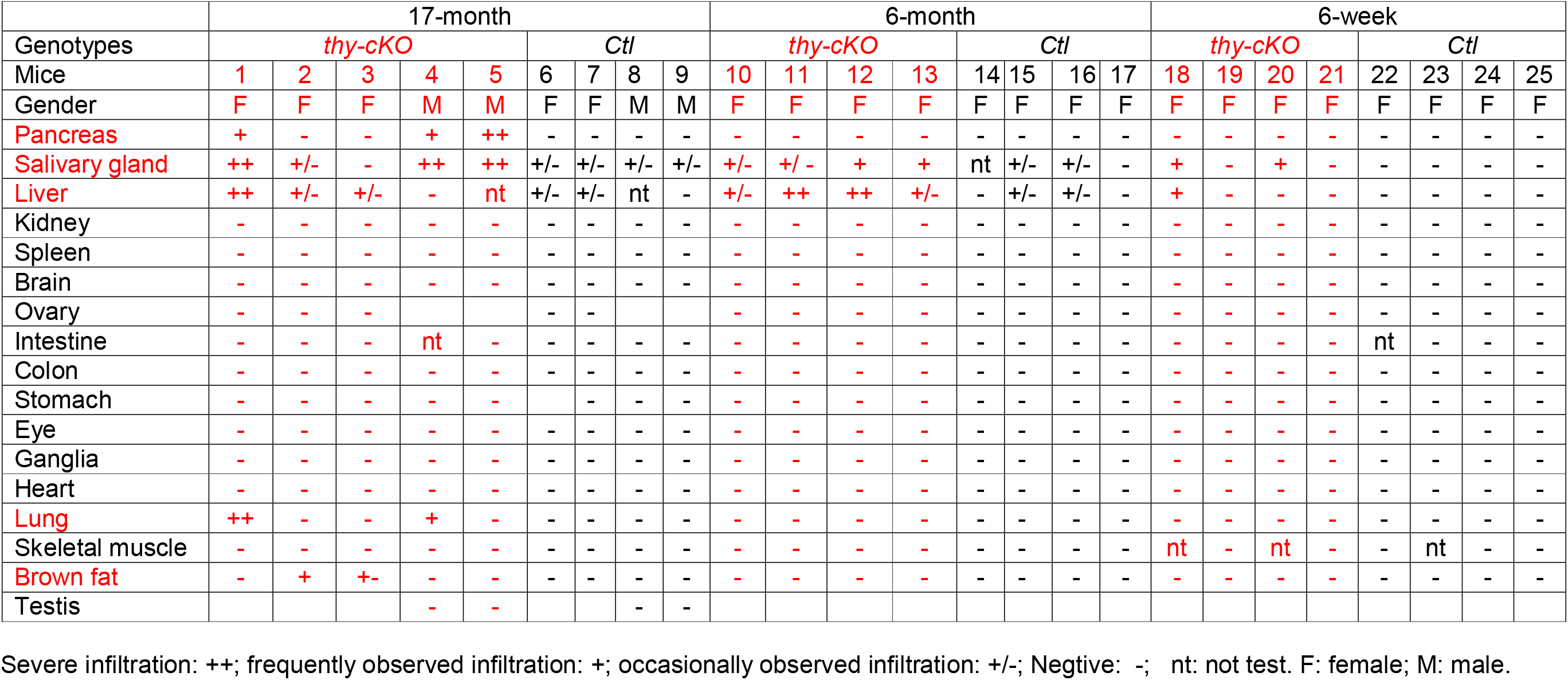
Lymphocytes infiltration in tissues.

**Table 2.**
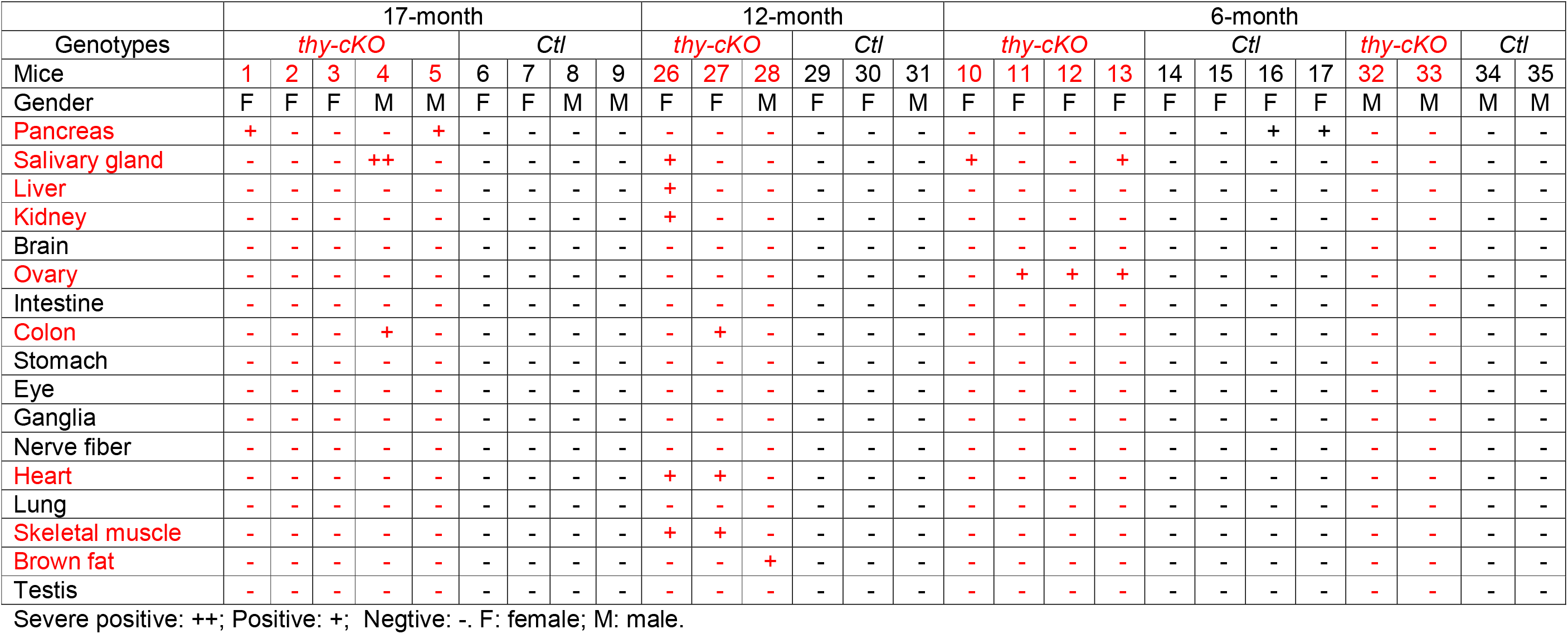
Autoimmune antibody reaction in tissues.

## Discussion

Insm1 was previously described as a factor functioning in neuroendocrine cells and neurons (27–29, 36–38). Herzig, Y *et al* reported the first evidence indicating Insm1 is a candidate transcriptional regulator of *Aire* (25). However, to our knowledge functional data on a role of Insm1 in thymus have not been systematic analyzed. We used *Insm1* null mutant mice, thymus-specific *Insm1* mutants and transplanted the thymus of *Insm1* mutants for an analysis of a role of Insm1 in autoimmunity.

We detected that *Insm1* is expressed in mTECs. Majority of Insm1 is expressed in Aire positive cells while a minor population of Insm1 positive cells did not express Aire. Using the published scRNA-seq data that were generated from post-Aire mTECs (18), we found that *Insm1* is expressed in post-Aire neuroendocrine mimetic cells. In *Insm1* mutant mTECs, the expression of *Aire*, the Aire-dependent and -independent TRAs were decreased. In consistent with the gene expression, *Insm1* mutation results in less Aire-positive cells in developmental mTECs but not the adult mTEC. Considering the cellular difference between perinatal and adult, i.e., Aire-expressing mTEC are enriched in post-Aire mTECs at the perinatal stage (18), Insm1 may contribution to the development of fetal Aire-expressing mTEC and the establishment of tolerization. To identify the regulatory role of Insm1 in mimetic cells, we examined the mimetic cell-type-specific gene expression and found that only those genes expressed in the neuroendocrine, enterohepatic, and pancreatic cells was downregulated in *Insm1* mutation. As Insm1 is co-expressed with Aire and consistently expressed only in post-Aire neuroendocrine mimetic cells, the alteration of gene expression in enterohepatic and Ptf1a^+^pancreatic cells may reflect the developmental regulation of Insm1 during the differentiation process of Aire-expressing mTECs into enterohepatic and Ptf1a^+^pancreatic cells. When *Insm1* overexpression was induced in the thymus, enlarged areas of Krt5 expression were observed, and the expression levels of *Aire* and TRA genes were increased. Furthermore, neuroendocrine cell marker genes were induced. The data further confirm the regulatory role of Insm1 in development of Aire-expressing mTEC and the post-Aire neuroendocrine mimetic cells.

The molecular and cellular model of TRA expression showed that Aire-expressing mTEC further differentiated into mimetic cells and both of Aire-expressing mTECs and mimetic cells are essential for the establishment of self-tolerance (23). In our study, we showed that *Insm1* is expressed in Aire-expressing mTECs and neuroendocrine mimetic cells, indicating that Insm1 is expressed in serial developmental stages of mTECs, i.e., widely expressed in Aire-expressing mTECs and restricted in neuroendocrine mimetic cells. Mutation of Insm1 impaired the gene expression in both Aire-expressing mTEC and neuroendocrine cells. The deficits of gene expression in Aire-expressing mTECs may contribute to the alteration of the gene expression in mimetic cells, such as the enterohepatic cells and Ptf1a^+^ pancreatic cells. Thus, Insm1 contributes to the development and function of Aire-expressing mTECs and post-Aire neuroendocrine mimetic cells.

Interestingly, we observed that Insm1 binding on the super-enhancer regions in mTECs. These super-enhancers contribute to the expression of *Insm1*-dependent genes. In addition, the majority of Insm1 binding sites overlapped with those of Aire, pointing towards the functional importance of the set of cis-elements binding by both, Insm1 and Aire in mTEC. Although a high proportion of common Insm1 binding sites were observed in mTECs of E18.5 and adult mice, the dysregulated genes were not conserved between E18.5 and adult in *Insm1* mutants. Rather, more up-regulated genes were identified in *Insm1* mutant adult mTECs than that of E18.5. In addition, the Insm1-binding super-enhancers were significant correlated with the Insm1-dependent genes expressed at E18.5 but not at adult stages, one explanation is that Aire-expressing mTEC is more enriched in post-Aire mTECs in perinatal than adult stage, Insm1 may regulate gene expression mainly in Aire-expressing mTECs through binding of super-enhancers. As *Insm1* is also expressed in neuroendocrine mTECs, further analysis of Insm1 function in the individual sub-population is needed, particularly in neuroendocrine mimetic cells.

Mutation of *Insm1* results in about 50% decrease of *Aire* expression. A dosage effect of Aire in TRAs regulation was discussed controversially. *Herzig* et al showed that 50% decreased Aire expression in *Aire^+/-^* mice did not alter the Aire-dependent TRAs expression (25). In contrast, others demonstrated an 80-90% reduction in the expression of a particular TRAs (*Ins2, Tff3, Mup1* and *Spt1*) in *Aire^+/-^* compared to wild-type littermate control mice (39). Additional studies also showed that TRAs levels were regulated by dosage of *Aire* expression (40, 41). In the *Insm1* mutant thymus, we observed downregulated expression of Aire-dependent TRA genes, possibly due to the decreased Aire expression.

In nude mice transplanted with *Insm1* mutant thymuses or in thymus specific *Insm1* mutant mice, we observed lymphocyte infiltration and autoimmune antibodies. These autoimmune responses were directed against various tissues, among them are pancreatic islets, the salivary gland, the lung and the kidney. However, not all these were equally attacked by immune reactions. Whereas pancreatic islets and the salivary gland were affected frequently in both, nude mice transplanted with *Insm1* mutant thymuses or thymus specific *Insm1* mutant mice, immune reactions in other tissues were observed only in occasionally. In addition, autoimmune responses in brown adipose tissues were observed in *Insm1* mutants, and have not been reported for *Aire* mutant mice (4, 5, 9, 10, 12, 14). In conclusion, out data show that Insm1 is a novel regulator of mTEC development, gene expression and particularly of TRAs expression in mTECs. Mutation of *Insm1* can cause autoimmune reactions.

## Materials and Methods

### Animals

*Insm1^lacZ/lacZ^* mice were described (26). E18.5 embryos were collected, genotyped and analyzed. A thymus specific mutation of *Insm1* was introduced by crossing *Insm1^flox/flox^* (28) and *Foxn1Cre* mice (JAX No. 018448). Tissues collected from *Rag1^-/-^* mice (GemPharmatech No.T004753) were used for autoimmune antibody test. The *Insm1KI* mouse strain was generated by introducing Insm1 coding sequences preceded 5’ by a stop cassette, that can be removed by Cre, into the *Rosa26* locus (Cyagen, Suzhou, China) (Fig S8). The littermate wildtype or heterozygous animals were used for control animals in the analysis. All animal experiments were approved by the institutional animal care and use committee of the Jinan University (IACUC-20211123-02).

### Thymus transplantation

Thymuses were isolated from E18.5 *Insm1^+/lacZ^* and *Insm1^lacZ/lacZ^* mice and cultured for eight days in RPMI-1640 containing 10% FBS and 1.25 mM 2’-deoxyguanosine. One thymus was transplanted into renal capsule of a nude mouse (6-week-old, CD1 background) using the procedure described previously (35). Eight weeks after transplantation, the thymus and newly generated lymphocytes were collected from the nude mice, various tissues of the animals were also collected and analyzed.

### Thymic cell dissociation and mTEC isolation

The fetal and adult thymic cells were dissociated using Liberase (42). In brief, the thymuses were separated and connective tissues and fat were removed. For E18.5 animals, 2-3 thymuses with of same genotype were pooled, and thymic lobes were cut into small pieces in RPMI to release the lymphocytes. Lymphocytes were collected in those cases when the entire thymic cell population was analyzed. Otherwise, the thymuses were digested in 100ul RPMI1640 containing 0.1mg/ml Liberase (Roche, Germany) and 20U/ml DNase I at 37 °C for 4 minutes. The digestion procedure was repeated twice, and cells were collected after each round of digestion.

mTECs isolation was performed using a magnetic beads-based method as previously described (43). In brief, 1μl biotinylated UEA1 (Vector Laboratories, B-1065-2) was incubated with 25μl Biotin Binder Dynabeads (Thermo Fisher Scientific, 11047) for 30 minutes at room temperature. Unbound UEA1 was removed by washing in 1ml FACS buffer (PBS with 0.02%BSA and 5mM EDTA). Thymuses from 2-3 E18.5 fetal or one adult mice were incubated with above prepared 25μl UEA1-coated beads at 4°C for 30 minutes. Unbound cells were removed by three washes with FACS buffer and by additional incubation with CD45 S-pluriBeads (pluriSelect Life Science, SKU#70-50010-11), before mTECs was collected for analysis. The isolated mTECs were used for RNA-seq, Cut&Tag and qRT-PCR analysis.

### Flow cytometry analysis

For analysis of lymphocytes in the transplanted thymuses, thymic lobes were gentle cut, released lymphocytes were collected and directly stained with CD4 and CD8 antibodies in FACS buffer for flow cytometric analysis. For analysis of thymic cells, stroma cells were dissociated using Liberase as described above. Staining with antibodies recognizing cell surface proteins were performed directly in FACS buffer, whereas staining with antibodies recognizing nuclear antigens was performed after fixation for 10 minutes in 4% PFA. Foxp3 staining was performed in Foxp3/Transcription Factor Staining Buffer (Invitrogen, 00-5523-00). Immunostaining procedures were performed as described (44). The antibodies used were listed in the supplementary table 5.

BD FACSCanto II or FACSAria II were used to analyze and collect cells (Beckton Dickenson, Franklin Lakes, USA). Cell quantification was determined using FloJo 7.6.5 software.

### Immunohistochemistry, western blot analysis and hematoxylin and eosin (H&E) staining

Immunohistochemistry and western blot analysis were performed as described (28). Fluorescence was imaged on a Zeiss LSM 700 confocal or a Leica DMi8 microscope, and the images were processed using Adobe Photoshop software. The antibodies used are listed in supplementary table 5. For the autoimmune antibody tests, serum of the *KO/nu* or *Thy-cKO* mice was used at the concentration of 1:25 on tissue sections obtained from *Rag1^-/-^* mice.

For H&E staining, the Lillie-Mayer method was used. In brief, 5% aluminum ammonium sulphate and 0.5% hematoxylin were used to stain the nuclei, followed by 0.3% acid alcohol incubation for the differentiation of nuclear staining. Acetified eosin was used for counterstain to reveal cellular detail.

### *Insm1* mutants RNA-seq analysis

For E18.5 mice, six to nine thymuses of each genotype (*Insm1* mutant and wildtype) were collected, and mTECs were isolated after pooling two to three thymuses of the same genotype. For 6-8 weeks mice, mTECs were isolated from each of the thymus for each genotype. Total RNAs were isolated using TRIZOL reagents. Three independent sequencing libraries for each genotype were generated using the NEBNext Ultra RNA Library Prep Kit for Illumina (NEB, USA). Illumina NovaSeq 6000 was used for sequencing, and 150-nt paired-end reads were generated. Sequencing data (6 Gb or more) with more than 94% of bases scoring above Q30 (accuracy rate 99.9%) were produced from each library sample. RNA-seq fastq files were aligned to the mouse genome (mm10) using STAR (v2.5.2a) (45) with parameter --outFilterMismatchNmax 2. The proportions of uniquely mapped reads were 79.4%-85.9%. Gene read numbers were counted using HTSeq (v0.11.0) (46) based on mouse gene annotation Ensembl v79, and the value of fragment per kilobase of exon per million reads mapped (FPKM) was calculated. Next, we used DESeq2 (v1.34.0) (47) to detect dysregulated genes under cutoff FDR≤0.1 and fold change≥1.5. To analyze downregulated TRAs and dysregulating-trend genes for down-steam analysis, a looser cutoff (*p*-value≤0.05 and fold change≥1.5) was performed. All the RNA-seq data are accessible on Gene Expression Omnibus (GSE193929).

For comparison of genes dysregulated after *Aire* mutation, published data were used (31). Gene read counts were downloaded from GEO (ID:GSE144877). Differential expressed genes were analyzed according to the method used in the referred paper (31).

GO enrichment analysis was performed by R package topGO (v2.38.1) (48) with “classic” algorithm and “Fisher exact test”, followed by the Benjamini-Hochberg multiple test correction using FDR=0.05 as a cutoff.

### qRT-PCR analysis

Cells were lysed and total RNA was isolated using TRIZOL reagent (Invitrogen). For qRT-PCR analysis, RNA was isolated from thymuses or mTECs of E18.5 fetal mice. cDNA was synthetized using HiScript II Q RT SuperMix for qPCR (+gDNA wiper) (R223-01, Vazyme, China) and analyzed using SYBR Green I based real-time quantitative PCR method on a CFX96 RT-PCR system (Bio-Rad). Expression levels were determined using the 2^-Δ Δ Ct^ method and *ActB* or *Krt5* as internal standards, and displayed as proportion of control. Primers used for quantitative analysis are listed in supplementary table S6.

### Insm1 CUT&Tag-seq analysis

CUT&Tag (Cleavage Under Targets and Tagmentation) analysis was performed according to the instruction of the manufacturer (Novoprotein, N259-YH01) and previously published protocols (49). In brief, mTECs were isolated using UEA1 coupled Dynabeads from wildtype mice. 0.5μl of Insm1 antibody or 1μg IgG antibody was incubated overnight at 4°C with 5×10^4^ mTEC cells in a volume of 100μl first antibody buffer. Donkey anti-guinea pig (Jackson ImmunoResearch,706-005-148) secondary antibody was used and incubated with the mTEC for 45 minutes in 100μl secondary antibody buffer at room temperature. After washing to remove unbound antibodies, the ChiTag transposon was incubated with mTECs for one hour at room temperature. Fragmentation, DNA purification and library construction were performed according to the protocol of the manufacturer. At least three biological replicas were obtained.

Illumina NovaSeq 6000 was used for sequencing, and 150-nt paired-end reads were generated. At least twelve Gigabyte pass filtering sequencing data was generated. Raw CUT&Tag sequencing data was trimmed to remove adaptor sequences using fastp (v0.12.0) (50) and aligned to mouse genome (mm10) using Bowtie2 (v2.2.9) (51) with the parameters “--local --very-sensitive-local --no-unal --no-mixed --no-discordant --phred33 -X 700”. Peaks were called using MACS2 (v2.2.6) (52) with the parameter “--keep-dup auto”. IgG data were used as a control. Insm1-binding peaks identified in three replicates with q-value<10e-5 were merged for subsequent analysis.

To calculate the proportions of Insm1 binding on different genomic regions, peaks were assigned to different genomic regions based on their summit positions on the Ensembl transcriptome annotation according to the priority order of CDS, 5’-UTR, 3’-UTR, promoter, ncRNA, intron, and intergenic, where the promoter were defined as the regions of −2000 bp to +500 bp of transcription start site (TSS). The CUT&Tag sequencing data are accessible on Gene Expression Omnibus (GSE193929).

### Single-cell RNA sequencing data analysis

Preprocessed transcript-by-cell matrix of scRNA-seq data of Pdpn^-^CD104^-^ mTEC^lo^ from perinatal and adult mice was downloaded from GEO (ID:GSM5831744), and read by scanpy (53) for downstream analysis. First, the valid 8,236 mTECs used in the study (18) were selected according to the barcode list provided. The gene expression matrix was normalized to counts per 10,000 counts, followed by log-transfer and scaling to unit variance and zero mean. Next, the top 2,000 highly variable genes were identified and selected, and PCA was performed for dimensionality reduction. According to the elbow point of the PC contribution curve, the top 30 PCs were used for the neighborhood graph construction, and UMAP was created for visualization. Cell clustering were preformed using the Leiden method (resolution=1.9). The scores of gene sets were calculated using the scanpy function tl.score_genes.

### TRA gene analysis

the list of TRA genes were identified in an earlier study (54) and provided by the authors (Prof. Perreault C, Université de Montréal, personal communication). Aire-dependent TRAs were identified by the overlaps of downregulated genes from the public Aire mutant RNA-seq data (31). The TRA expression profiles of various mouse tissues were generated in pervious microarray studies (33) and downloaded from GEO (GSE10246). To visualize TRA expressions in different tissues (Figure S3), samples of each tissue were merged by mean expression values, followed by z-score normalization on each gene.

### Super enhancer analysis

Super enhancer regions were identified in the published study (13) and were provided by the author via email (Prof. Diane Mathis, Harvard Medical School, personal communication). The control regions used in Fig 4B were the sequences with the same length as the super enhancer and with an offset of the distance of the super enhancer length plus 200kb.

## Acknowledgments

The authors thank Prof. Carmen Birchmeier (Max-Delbrück-Center for Molecular Medicine,Germany) for helpful discussions at the initiation of the project, for providing the *Insm1^+/lacZ^ Insm1^+/flox^* animals and for critical reading of the manuscript. The authors thank Dr. St-Pierre, C (University of Montreal) and Prof. Perreault, C (University of Montreal) for kindly sharing the detailed information of TRAs. The authors thank Prof. Diane Mathis (Harvard Medical School) for providing the super-enhancer coordinates. The authors thank Prof. Yuanzhi Lu (Department of Pathology, The First affiliated hospital of Jinan University) for the help in analyzing lymphocytes infiltration.

## Funding

This work was supported by the National Natural Science Foundation of China (31970856) and the Clinical Frontier Technology Program of the First Affiliated Hospital of Jinan University, China (JNU1AF-CFTP-2022-a01236).

## Author Contributions

W.T. designed the study and performed the molecular experiments. Y.W. and J.W. contributed to the animal experiments, tissues analysis and flow cytometry analysis. Y.W., Z.Y., W.Y. and G.Y. contributed to the molecular experiments, cells and tissues experiments. J.X. performed and managed the bioinformatic analysis and participated manuscript preparation. S.J. supervised the project, analyzed the data and wrote the manuscript. S.J. is the guarantor of this work and takes responsibility for the integrity of the data and the accuracy of the data analysis.

## Notes

### Competing Interest Statement

The authors have declared no competing interest.

### Summary of Updates

section on result/discussion updated to clarify the Insm1 expressing and function in mTECs with related Fig1 and 3 revised

