## Supplementary figures and images for "Insm1 regulates the development of mTECs and immune tolerance"

### Fig S1

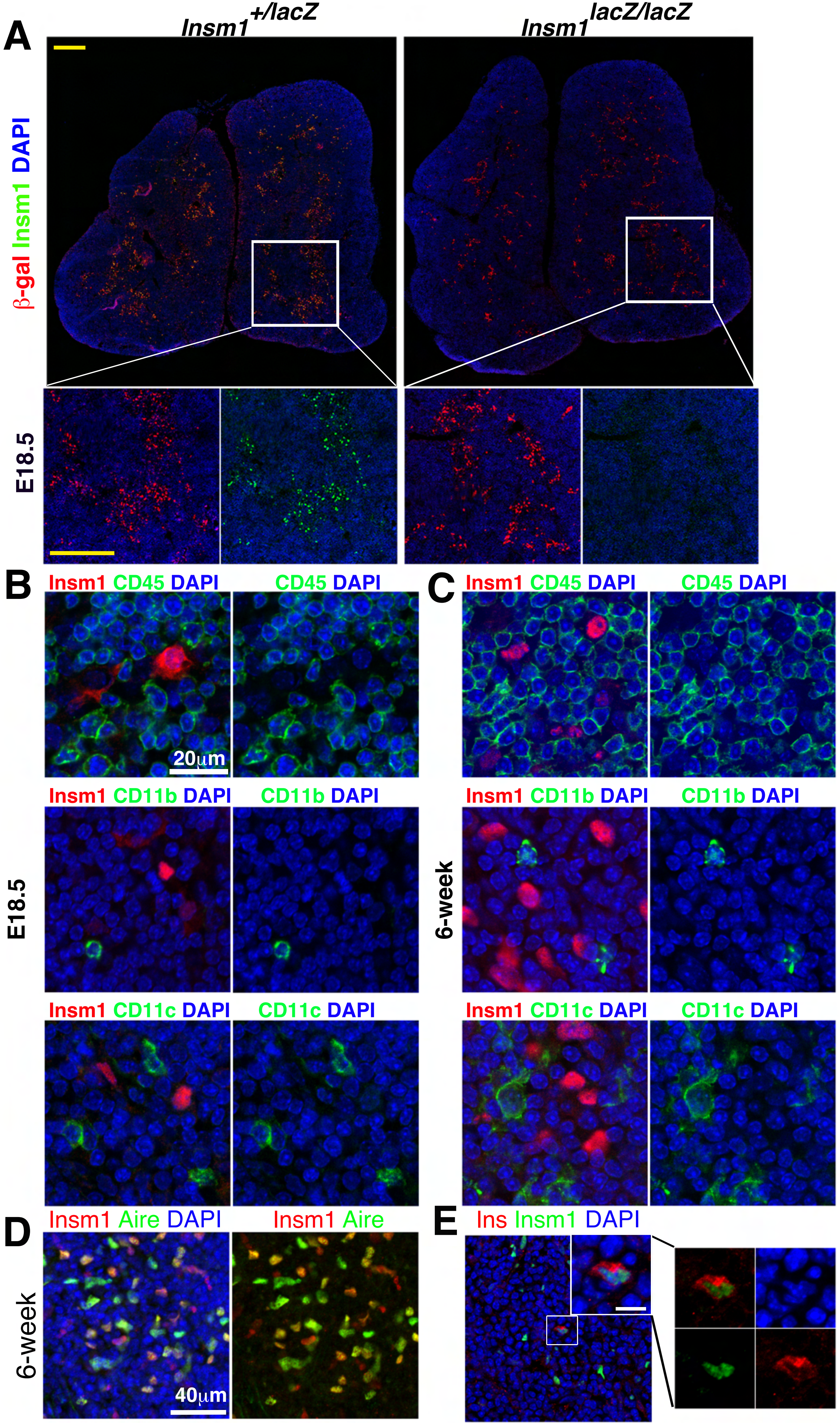

### Fig S2

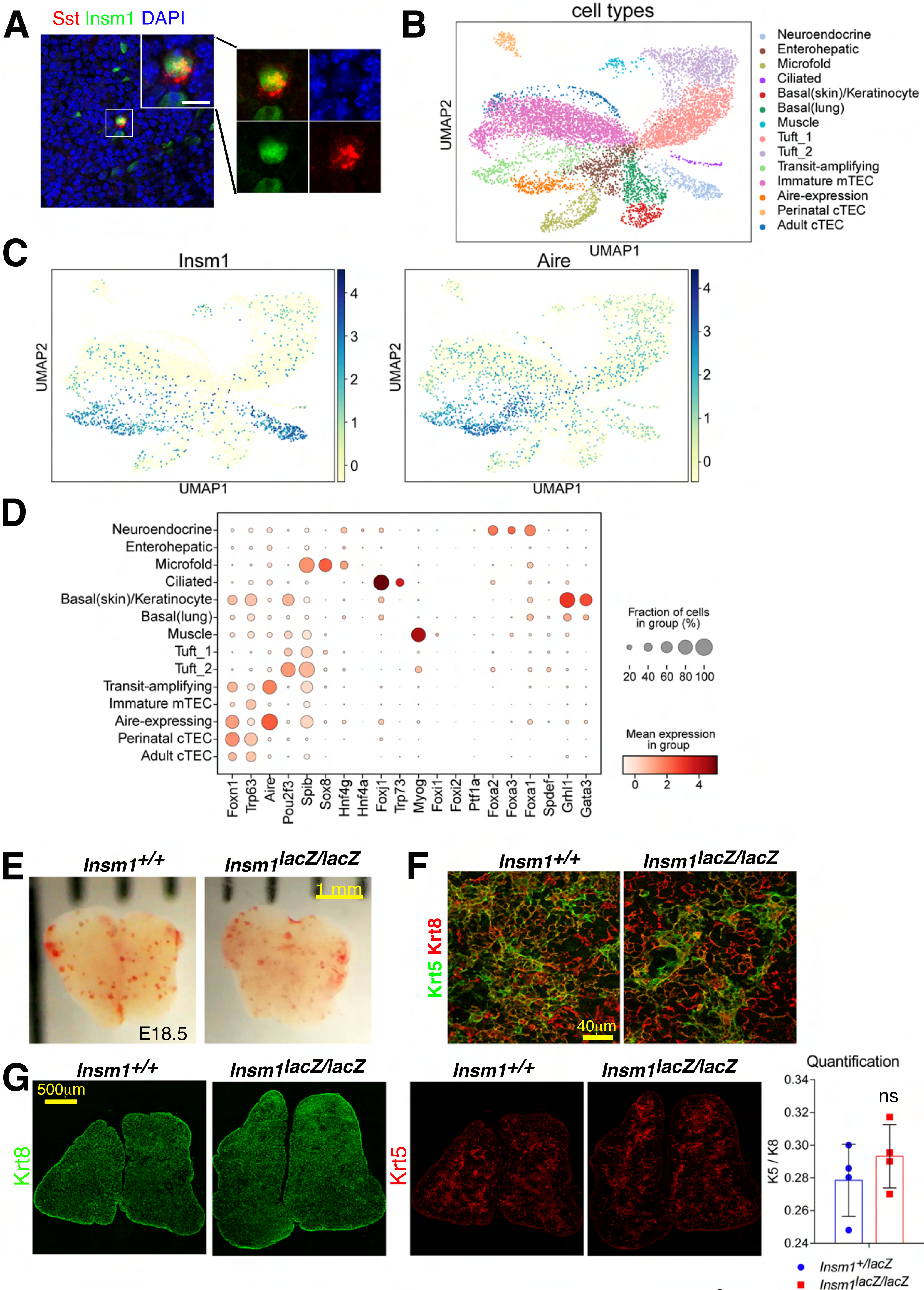

### Fig S3

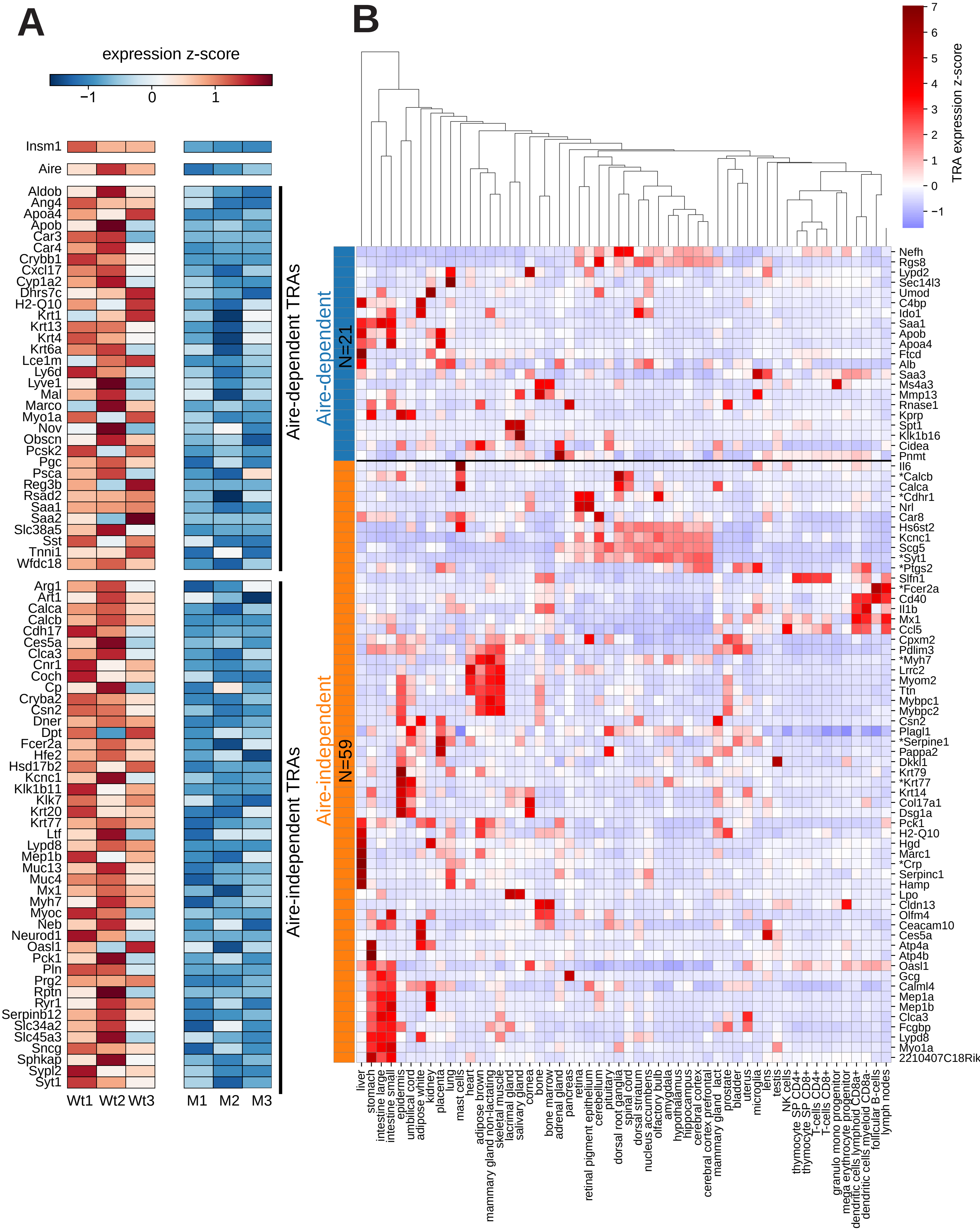

### Fig S4

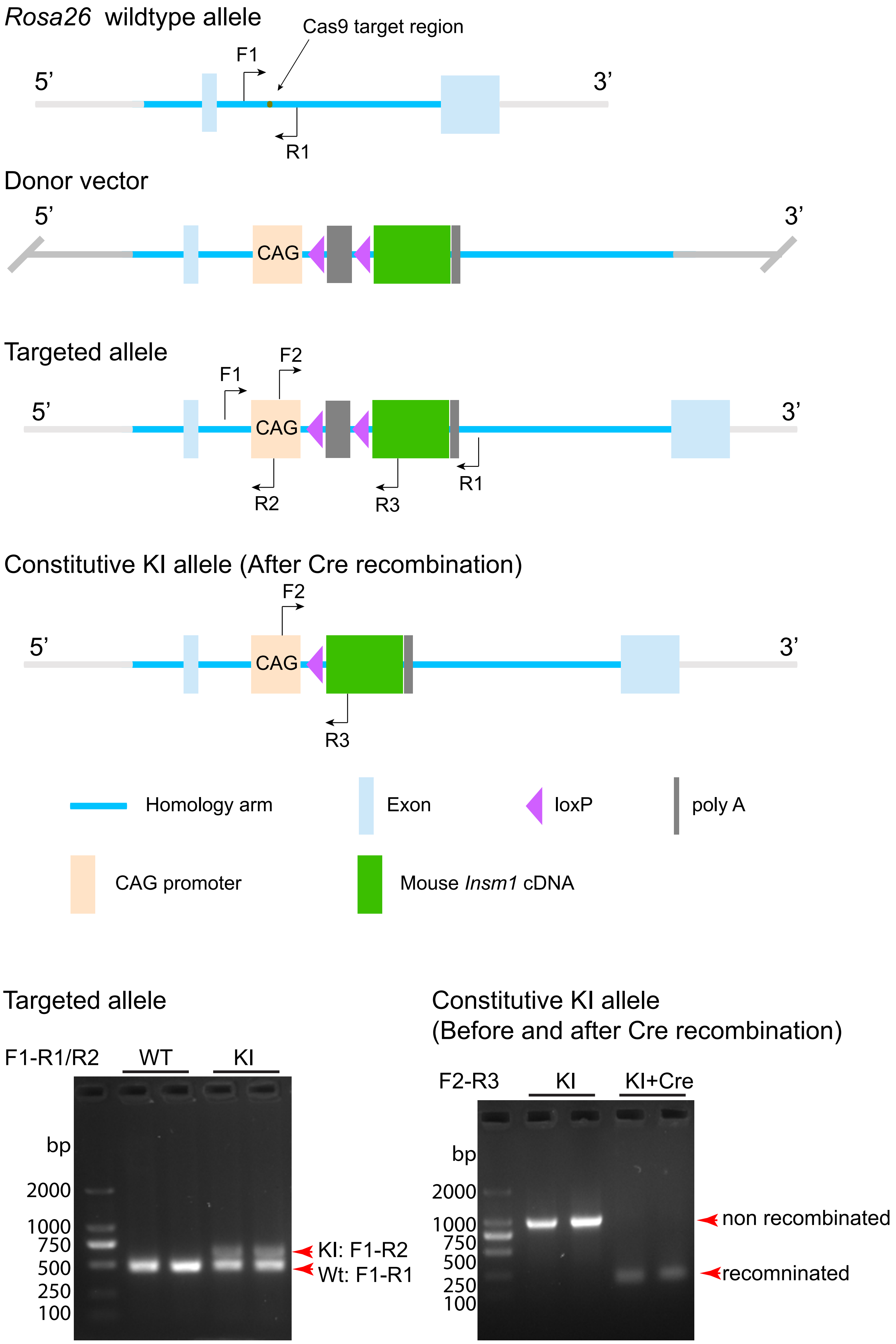

### Fig S5

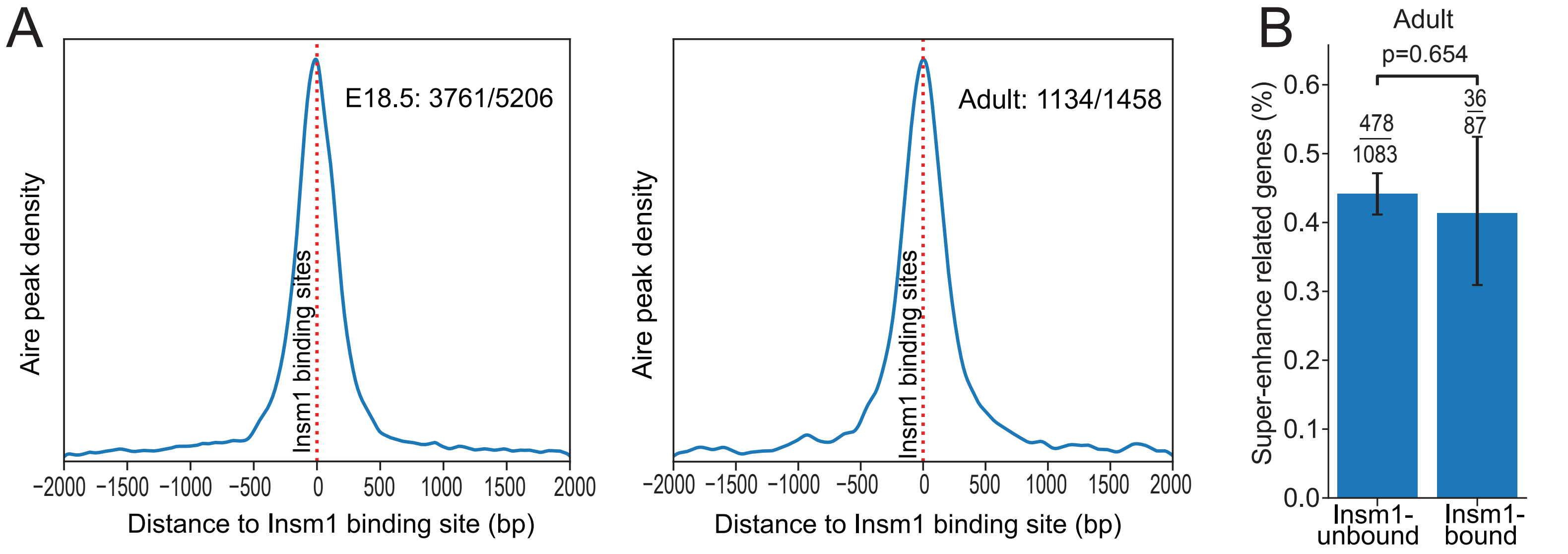

### Fig S6

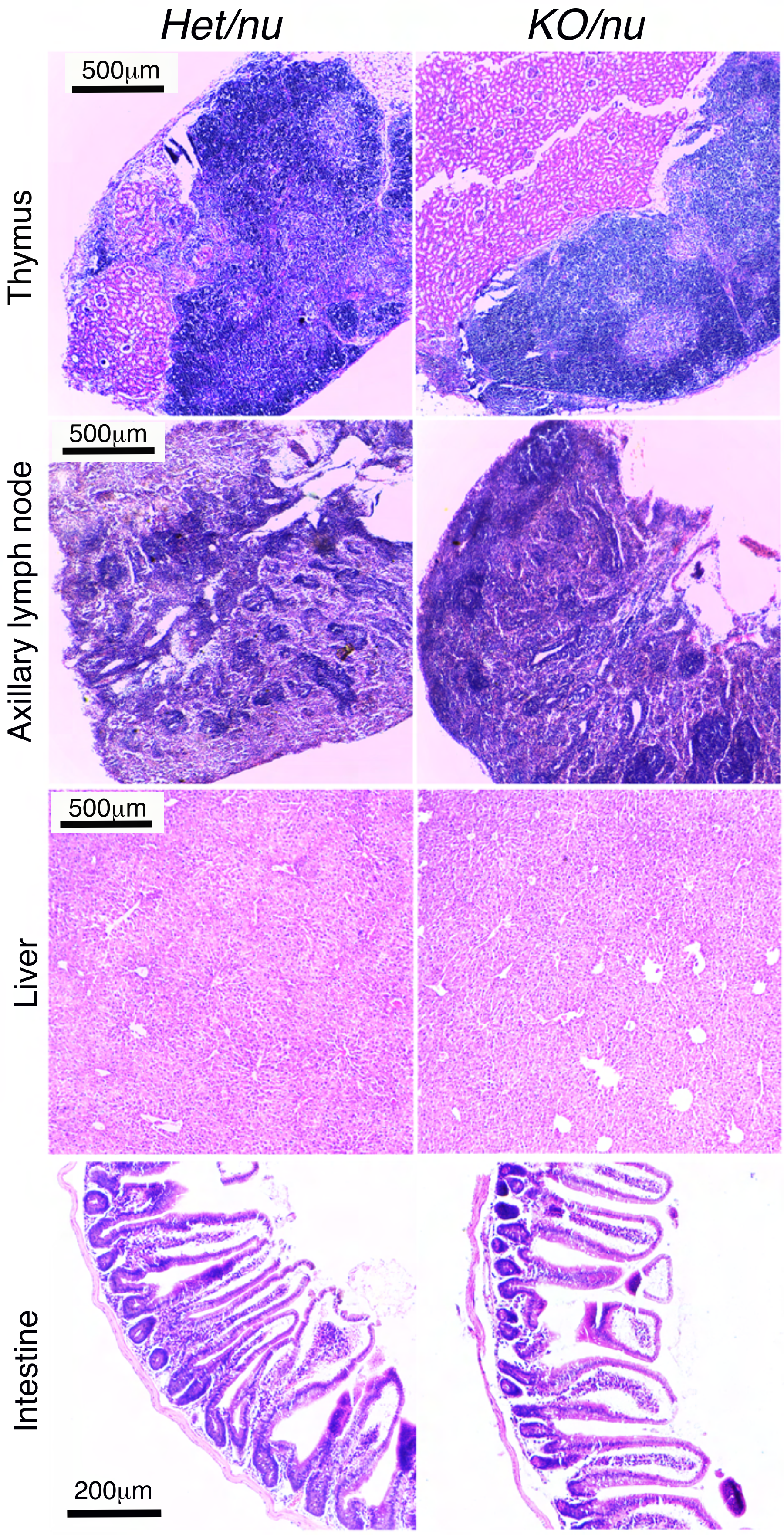

### Fig S7

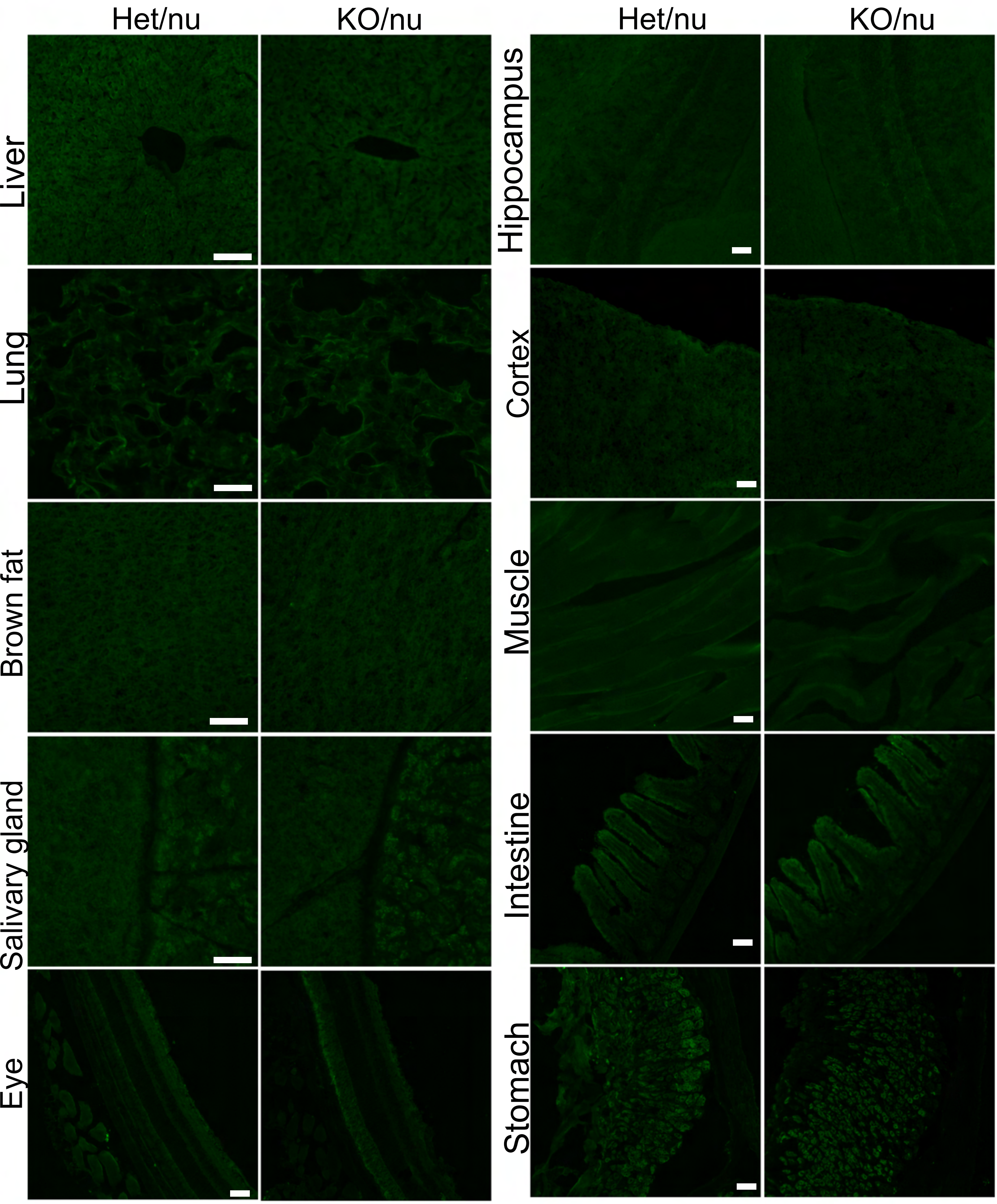
