## supplementary figure legends for "Insm1 regulates the development of mTECs and immune tolerance"

**Figure S1. Insm1 expression in the thymus**

(A) Immunofluorescence analysis of the thymus using antibodies against Insm1 and β-gal in thymus of *Insm1^+/lacZ^* and *Insm1^lacZ/lacZ^* mice. Shown are sections stained with anti-β-gal (red) and anti-Insm1 (green) antibodies. DAPI was used as a counterstain. The lower panels showed the magnification of the indicated areas. (B,C) Immunofluorescence analysis of Insm1 (red) and CD45, CD11b or CD11c (green, as indicated in the panels); analyzed are the thymus of E18.5 (B) and 6-week-old (C) mice. DAPI was used as a counterstain. (D) Immunofluorescence analysis of Insm1 (red) and Aire (green); analyzed is the thymus of 6-week-old mice. (E) Immunofluorescence analysis of thymuses of 6-week-old mice using antibodies against Insulin (red) and Insm1 (green). Scale bar = 40μm and 10μm in the main and magnified panels, respectively.

**Figure S2. Insm1-positive cell types in the thymus**.

(A) Immunofluorescence analysis of thymuses of 6-week-old mice using antibodies against (A) Sst (red) and Insm1 (green). Scale bar = 40μm and 10μm in the main and magnified panels, respectively. (B) the UMAP generated using the published data (1). Showing are the post-Aire mTECs Pdpn^-^CD104^-^mTEC^lo^ mTEC cells, which contains the mimetic cells. (C) the expression of Insm1 (left) and Aire (right) in sub-populations of the mTEC. (D) Expression of mimetic cell lineage transcription factors in the cell-clusters analyzed using the published post-Aire mTECs scRNAseq (1). (E) Isolated thymuses of wildtype littermate control and *Insm1* mutants at E18.5. (F-G) immunofluorescence analysis of Krt5 (green) and Krt8 (red) of the thymus of wildtype littermate control and *Insm1* mutants. (G right) Quantification of the ratio of Krt5 and Krt8 positive areas in thymuses of wildtype littermate control and *Insm1* mutants (n=4). Data are presented as means ± SD, statistical significance was assessed by 2-tailed unpaired Student's t-test. ns: *P>*0.05

**Figure S3. Gene expression analysis**

(A) Heatmap of dysregulated TRA genes in mTEC of E18.5 mice. (B) Heatmap showing the expression of downregulated TRAs in different mouse tissues using the public microarray data.

**Figure S4. Scheme of the generation of the *Insm1KI* allele**

The wildtype *Rosa26* locus was used to insert *Insm1* sequences between exon 1 and 2 by CRISPR-Cas9 based targeting. The donor vector contained a floxed 3xSTOP cassette upstream of the *Insm1* cDNA. The structure of the targeted allele, and the allele *after* Cre mediated recombination were verified by PCR using the indicated F1-R1/R2 and F3-R3 primer sets, respectively. The homology arm, exon sequences (light blue), *loxP* sequences (magenta), *polyA* sequences (dark gray), *CAG* promoter (light orange) and *Insm1* sequences (green) are indicated.

**Figure S5. Analysis of Insm1 binding sites.**

(A) Comparison of the position of Aire and Insm1 binding sites displayed as an density curve of Aire binding peaks aligned to the summit of the Insm1 binding peaks in E18.5 (left) and adult (right). Co-binding sites and total Insm1 binding sites were showed as numerators and denominators respectively on the right corner. (B) Proportions of super-enhancers located within ±500kb of genes that were dysregulated in *Insm1* mutant mTECs (p-value≦0.05, FC≧1.5) at the adult stage. Stratified by the super-enhancers bound or unbound by Insm1.

**Figure S6. Structure of tissues isolated from transplanted animals.**

H&E staining of the thymus, axillary lymph node, liver and intestine from nude mice transplanted with thymus of *Insm1^+/lacZ^* or *Insm1^lacZ/lacZ^* mice.

**Figure S7. Examples of the absence of autoimmune reactions in individual animals**

Immunofluorescence analysis using serum (green) isolated from nude mice transplanted with a thymus of *Insm1^+/lacZ^* or *Insm1^lacZ/lacZ^* mice. Analyzed are multiple tissues of *Rag1^-/-^* mice as indicated in the figure.

1.	Michelson DA, Hase K, Kaisho T, Benoist C, & Mathis D (2022) Thymic epithelial cells co-opt lineage-defining transcription factors to eliminate autoreactive T cells. *Cell* 185(14):2542-2558 e2518.
